# The Drosophila HP1 family is associated with active gene expression across chromatin contexts

**DOI:** 10.1101/2020.02.24.958538

**Authors:** John M. Schoelz, Justina X. Feng, Nicole C. Riddle

## Abstract

Drosophila Heterochromatin Protein 1a (HP1a) is essential for heterochromatin formation and is involved in transcriptional silencing. However, certain loci require HP1a to be transcribed properly. One model posits that HP1a acts as a transcriptional silencer within euchromatin while acting as an activator within heterochromatin. However, HP1a has been observed as an activator of a set of euchromatic genes. Therefore, it is not clear whether, or how, chromatin context informs the function of HP1 proteins. To understand the role of HP1 proteins in transcription, we examined the genome-wide binding profile of HP1a as well as two other Drosophila HP1 family members, HP1B and HP1C, to determine whether coordinated binding of these proteins is associated with specific transcriptional outcomes. We found that HP1 proteins share a majority of their endogenous binding targets. These genes are marked by active histone modifications and are expressed at higher levels than non-target genes in both heterochromatin and euchromatin. In addition, HP1 binding targets displayed increased RNA polymerase pausing compared to non-target genes. Specifically, co-localization of HP1B and HP1C was associated with the highest levels of polymerase pausing and gene expression. Analysis of HP1 null mutants suggests these proteins coordinate activity at transcription start sites (TSSs) to regulate transcription. Depletion of HP1B or HP1C alters expression of protein-coding genes bound by HP1 family members. Our data broadens understanding of the mechanism of transcriptional activation by HP1a and highlights the need to consider particular protein-protein interactions, rather than broader chromatin context, to predict impacts of HP1 at TSSs.

## INTRODUCTION

Non-histone chromosomal proteins are essential to ensure genome integrity and function (Filion *et al*. 2010; Kharchenko *et al*. 2011). One prominent class of non-histone chromosomal proteins is represented by the Heterochromatin Protein 1 (HP1) family (Vermaak and Malik 2009; Canzio *et al*. 2014; Eissenberg and Elgin 2014). HP1 proteins are characterized by their unique domain structure consisting of a chromo-domain and a chromoshadow-domain connected by a hinge region (Smothers and Henikoff 2001). The chromo-domain mediates interactions between HP1 proteins and methylated histone tails (Jacobs *et al*. 2001), while the chromoshadow-domain mediates HP1 protein dimerization and interactions between HP1 family members and proteins containing a PxVxL amino acid motif (Thiru *et al*. 2004; Lechner *et al*. 2005). The ability to bind both methylated histones and a diverse set of additional nuclear proteins confers the classification of ‘hub protein’ to the HP1 family. As such, HP1 proteins are active in several different nuclear processes including heterochromatin formation (Larson *et al*. 2017; Strom *et al*. 2017; Machida *et al*. 2018), DNA repair (Ryu *et al*. 2015; Amaral *et al*. 2017), DNA replication (Li *et al*. 2011), and regulation of gene expression (Danzer and Wallrath 2004; Lin *et al*. 2008; Kwon *et al*. 2010), illustrating the importance of this gene family (Badugu *et al*. 2003; Vermaak and Malik 2009).

The *Drosophila melanogaster* HP1 family includes five full-length genes (containing both a chromo-domain and a chromoshadow-domain): *Su(var)205* (encoding the HP1a protein), *HP1b*, *HP1c*, *rhino* (encoding HP1D), and *HP1e* (Vermaak and Malik 2009). *Su(var)205*, *HP1b* and *HP1c* are expressed ubiquitously while *rhino* and *HP1e* are present mostly in female and male germ cells, respectively (Vermaak *et al*. 2005; Levine *et al*. 2012). Based initially on studies from Drosophila polytene chromosomes, the HP1a protein mostly localizes to pericentric heterochromatin, telomeres, chromosome four, and a few euchromatic loci (James *et al*. 1989; Fanti *et al*. 2003). This localization pattern was confirmed by later chromatin immunoprecipitation (ChIP) studies from the modENCODE (model organism encyclopedia of DNA elements) consortium and others (Riddle *et al*. 2011; Ho *et al*. 2014). HP1B localizes throughout heterochromatic and euchromatic domains on polytene chromosomes, and HP1C localizes mostly to euchromatin (Smothers and Henikoff 2001). These patterns are reinforced also by data from ChIP-chip and ChIP-seq experiments performed by the modENCODE consortium and others (Ho *et al*. 2014). Loss of function mutations in the *Su(var)205* gene encoding HP1a disrupt the formation of heterochromatin and are homozygous lethal (Eissenberg *et al*. 1990), while loss of function mutations in the *HP1b* and *HP1c* genes are homozygous viable (Font-Burgada *et al*. 2008; Mills *et al*. 2018). This finding has led to the speculation that the HP1B and HP1C proteins may exhibit functional redundancy. Together, these data provide a model of the Drosophila HP1 family wherein HP1a is an essential heterochromatin protein, HP1C is a non-essential euchromatin protein, and HP1B is a non-essential protein binding to both heterochromatin and euchromatin. This model is a starting point for investigating the individual roles of HP1 family proteins in diverse biological processes.

While the role of the Drosophila HP1 family in the regulation of gene expression is complex, HP1a is well-known for its role in the formation of heterochromatin, associating it with transcriptional silencing activity (James *et al*. 1989; Eissenberg *et al*. 1990; Li *et al*. 2003; Danzer and Wallrath 2004). Here, HP1a recognizes and binds histone three, lysine nine di- and trimethylation (H3K9me2/3) through its chromo-domain and subsequently recruits the H3K9 methyltransferase Su(var)3-9, resulting in the propagation of heterochromatic domains and the silencing of transposable elements (TEs) (Czermin *et al*. 2001; Jacobs *et al*. 2001; Snowden *et al*. 2002; Motamedi *et al*. 2008). HP1a is also critical for the establishment and maintenance of a phase separated environment between heterochromatin and euchromatin which is thought to limit contact between transcriptional activators and the heterochromatic compartment of the genome (Larson *et al*. 2017; Strom *et al*. 2017; Sanulli *et al*. 2019). These observations lead to a model of HP1a functioning as a transcriptional repressor, which is supported by data from studies tethering HP1a to transgene reporters that result in transcriptional silencing (Li *et al*. 2003; Danzer and Wallrath 2004). Complicating this model, however, is the observation that a number of both euchromatic and heterochromatic loci require HP1a to maintain an active transcriptional state (Lu *et al*. 2000; Cryderman *et al*. 2005). Additionally, inducible loci such as heat shock response genes are enriched for HP1a upon induction (Piacentini *et al*. 2003; Piacentini *et al*. 2009). One proposed model to explain these differences is that HP1a serves different functions in different chromatin contexts through interactions with distinct sets of protein partners (Li *et al*. 2002). However, evidence for this hypothesis is lacking.

An alternative approach to investigating the effects of HP1a on gene expression is to focus on its interactions with other HP1 family proteins. While the exact function of HP1B or HP1C in transcriptional regulation is not well characterized, tethering studies of transgene reporters support a role for HP1C in transcriptional activation (Font-Burgada *et al*. 2008). Evidence for the impact of HP1B on gene transcription is conflicting. While tethering studies support a role for HP1B in gene silencing, PEV studies support a role for HP1B in transcriptional activation (Font-Burgada *et al*. 2008; Mills *et al*. 2018). HP1C recruits the Facilitates Chromatin Transcription (FACT) complex to promote RNA polymerase II (RPII) elongation after being targeted to chromatin by the zinc finger transcription factors WOC and ROW (Font-Burgada *et al*. 2008; Kwon *et al*. 2010). Both HP1a and HP1B also interact with subunits of FACT as well as WOC, but the nature of these interactions is uncharacterized (Kwon *et al*. 2010; Ryu *et al*. 2014). RNA-Seq experiments following RNAi knockdown of all three HP1 paralogs in Drosophila reveal evidence of both activating and silencing functions of HP1 proteins: both widespread up- and down-regulation of target genes are observed with a large number of misregulated genes being shared across knockdown conditions (Lee *et al*. 2013). These findings raise the possibility that HP1 proteins may coordinate their activity to regulate gene expression of a common transcriptional program.

Here, we explore whether combinatorial action and cooperative activity of multiple HP1 proteins at a single locus may predict differences in transcriptional activity at protein-coding genes with better accuracy than knowledge of the surrounding chromatin context. To achieve this goal, we integrate ChIP-Seq and RNA-Seq datasets to characterize the genomic distribution of each HP1 protein and to measure the association between each HP1 protein and transcriptional states genome-wide. We find active transcription at binding targets shared between multiple HP1 proteins across a variety of chromatin states. Furthermore, these targets exhibit signatures of RNA polymerase II promoter proximal pausing, providing evidence for a potential mechanism for transcriptional activation by HP1 proteins. Analysis of pausing in HP1 null mutants suggests coordinated activity between HP1 family members is important for proper gene expression. These findings suggest knowledge of locus-specific protein-protein interactions is more informative for predicting HP1 function at transcription start sites (TSSs) than knowledge of a broader chromatin context.

## RESULTS

### Drosophila HP1 proteins are enriched in heterochromatin, but also bind throughout euchromatin

In order to better understand the function of the Drosophila HP1 family in transcriptional regulation, we set out to identify endogenous targets for all three somatic HP1 family members in the Drosophila genome: HP1a, HP1B, and HP1C. We began by re-analyzing existing ChIP-Seq data sets for HP1a, HP1B and HP1C from third instar larvae generated by the modENCODE consortium to characterize the genome-wide distributions of these proteins (Figure 1A) (Ho *et al*. 2014). We verified significant enrichment of HP1a (blue track, outer circle) within pericentric heterochromatin and on chromosome four, observing 34% and 2% of HP1a enriched regions resided in these chromatin domains, respectively. However, despite this enrichment, a majority (63%) of HP1a enriched domains resided in euchromatin (Figures 1A and B). Binding behavior of HP1a did not appear to be consistent across different chromatin domains. We found average HP1a-enriched domain widths of 14.8 kb and 21.7 kb within pericentric heterochromatin and on chromosome four, respectively, which was greater than the average width of 2.8 kb observed within euchromatin (Figure 1C, p < 2.2e-16, Mann-Whitney test). Meanwhile, an even greater majority of HP1B (green track, middle circle) and HP1C (pink track, inner circle) peaks were located in euchromatin, 87% and 95%, respectively (Figures 1A and B). For HP1B, we detected a greater share of signal within heterochromatin than for HP1C: 11% of HP1B peaks were within heterochromatin as opposed to just 4% of HP1C peaks (Figures 1A and B). A similar share of HP1B and HP1C peaks mapped to chromosome 4 (1%). We found that the average HP1B enriched peaks width of 2.7 kb and 4.3 kb within heterochromatin and on chromosome four were significantly larger than the average width of 2.3 kb within euchromatin (Figure 1D, p < 2.2e-16, Mann-Whitney test). Finally, the average HP1C peak width of 4.5 kb and 3.5 kb within heterochromatin and on chromosome four was also larger than the average peak width of 2.6 kb found within euchromatin (Figure 1E, p < 0.02, 3e-4, Mann-Whitney test). Thus, while in the literature HP1a is often characterized as a heterochromatin protein and HP1C as a euchromatin protein, all three somatically expressed HP1 proteins in Drosophila are found throughout both chromatin compartments, although their binding behavior differs somewhat across compartments.

**Figure 1.**
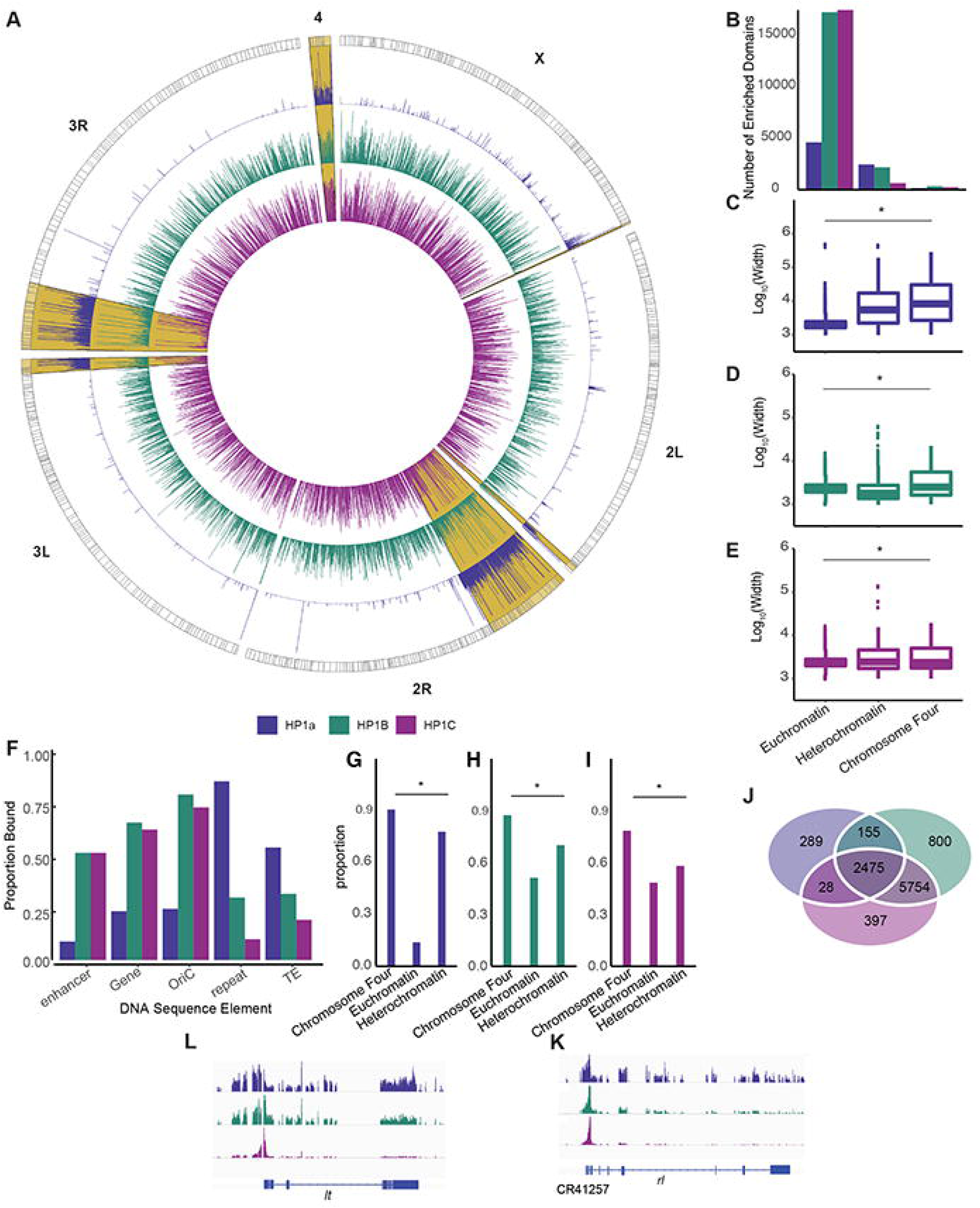
The genome-wide binding landscape of the Drosophila HP1 family. (A) Input-corrected genome-wide ChIP-Seq tracks for HP1a (blue track), HP1B (green track), and HP1C (purple track) in wildtype third instar larvae. Coverage is plotted as log_2_(ChIP/Input). Exterior gray track denotes positions of cytogenetic bands. Pericentric heterochromatic regions are highlighted in yellow using chromatin boundaries defined previously (Riddle *et al*. 2011). (B) Number of HP1-enriched domains identified in each chromatin compartment. (C-E) Comparison of enriched domain width across chromatin compartments for HP1a (C), HP1B (D), and HP1C (E). Average differences were evaluated using Mann-Whitney U test. (F) Comparison of HP1 protein occupancy at different DNA sequence elements. Y axis denotes the fraction of annotated elements bound by a given HP1 protein. (G-I) Fraction of genes bound by HP1 proteins in different chromatin compartments. Y axis denotes the proportion of genes bound by a given HP1 protein out of the total number of genes in that compartment. (J) HP1 proteins share a majority of their binding sites. The two most frequent combinations observed were co-localization of HP1B (green oval) and HP1C (purple oval) as well as both proteins colocalizing with HP1a (blue oval). (L-K) Genome browser screenshots of HP1 proteins colocalizing at heterochromatic genes *light* and *rolled*. Y axis denotes coverage. Gene structures are depicted beneath screenshots. (* denotes cutoff of p < 0.05)

To further examine the three HP1 proteins, we also looked at their tendency to localize to different DNA sequence elements. We investigated HP1 protein binding behavior at five different classes of DNA elements annotated in the Drosophila genome assembly (release dmel r6.25): enhancers, genes, origins of replication (OriCs), repeat regions, and TEs. For each DNA element, we measured the proportion of elements that overlapped with the binding site of an HP1 protein. HP1a bound the largest fraction of repeats and TEs among the three HP1 proteins, occupying approximately 80% and 50% of these elements respectively (Figure 1F), in agreement with a function for HP1a in TE regulation and silencing. In contrast, HP1B occupied approximately 25% of all TEs and repeats, while HP1C occupied approximately 20% of all TEs and was largely absent from repeat regions – HP1C is present at less than 10% of all repeat regions in the Drosophila genome (Figure 1F). Interestingly, OriCs marked a stark difference in HP1 binding behavior for the three proteins examined. HP1B and HP1C were present at approximately 75% of all OriCs, while HP1a was present at less than 25%. A similar trend was observed at genes and enhancers. HP1B and HP1C occupied greater than 50% of all protein coding genes and enhancers while HP1a occupied fewer than 25% of all genes and fewer than 5% of all enhancers. Thus, HP1 proteins can be differentiated by their tendency to localize to different DNA sequence elements, although those tendencies are not absolute.

### The HP1 family share genic binding sites in various chromatin contexts

Next, we set out to create a comprehensive list of HP1 binding targets across different chromatin contexts to quantify the extent to which HP1 proteins share binding sites at protein-coding genes. Within third instar larvae, HP1a accumulated to high levels at protein-coding genes located within heterochromatin and on chromosome four (Figure 1G). We found that 85% of all heterochromatic genes and 98% of all chromosome four genes were bound by HP1a. Meanwhile, HP1a was only present at 15% of euchromatic genes (Figure 1G, p < 2.2e-16). HP1B and HP1C were present at a high number of genes within heterochromatin and on chromosome four as well, but also occupied a larger share of euchromatic genes than HP1a (Figures 1H-I). HP1B was present at 70% of heterochromatic genes, 87% of chromosome four genes, and 50% of euchromatic genes (Figure 1H). HP1C bound 58% of heterochromatic genes, 78% of chromosome four genes, and 48% of euchromatic genes (Figure 1I). While all three HP1 proteins are enriched at genes located within heterochromatin and on chromosome four, they still bind a large number of euchromatic genes.

An overlap analysis of binding targets for all three HP1 proteins demonstrated that HP1 proteins share a majority of their binding sites in third instar larvae (Figure 1J & Supplemental Figure 2A). Two combinations of HP1 proteins were particularly widespread. HP1B and HP1C shared 90% of their binding sites at protein coding genes within third instar larvae and bound 32% of all protein coding genes. Meanwhile, all three HP1 proteins shared 13% of all bound genes (Figure 1J). This overlap can be illustrated by looking at two well-studied heterochromatin genes, *light* and *rolled* (Figures 1K-L). The significance of co-localization of HP1 family members is not understood, but it has been suggested previously that HP1 family proteins may display some degree of functional compensation (Ryu *et al*. 2014), particularly between HP1B and HP1C. All three HP1 proteins co-immunoprecipitate as well as form dimers through the chromoshadow-domain (Lee *et al*. 2019). It is unknown how these interactions affect gene expression.

### HP1 binding targets are highly expressed

To gain additional insights into the functions of the HP1 proteins in gene regulation, we characterized the protein-coding genes bound by HP1 proteins. We compared levels of expression between HP1 target and non-target genes using publicly available RNA-Seq data from third instar larvae (Supplemental Figure 1) (Mills *et al*. 2018). HP1a, HP1B, and HP1C target genes all exhibited significantly higher levels of expression than non-target genes (Supplemental Figures 1A, B, and C, p < 0.0002, permutation tests). High levels of expression at endogenous HP1 targets are surprising given the results of tethering studies which show both HP1a and HP1B act as transcriptional repressors at reporter genes (Font-Burgada *et al*. 2008).

Next, we performed gene ontology (GO) analysis (Huang da *et al*. 2009a; Huang da *et al*. 2009b) to further characterize endogenous HP1 binding targets, focusing on the biological process category of GO terms. Among HP1 binding targets, we identified significant enrichment for terms related to nervous system development and function such as ‘Response to axon injury’ (HP1a, Supplemental Figure 1B) and ‘Axon extension’ as well as ‘Synaptic vesicle coating’ (HP1B, Supplemental Figure 1D). Additionally, we observed terms broadly associated with mitosis and chromosome segregation including ‘Synaptonemal complex organization’ among HP1a binding targets (Supplemental Figure 1D), ‘Positive regulation of growth’ among HP1B binding targets, and ‘Spindle organization’ as well as ‘G1/S transition of mitotic cell cycle’ among HP1C binding targets (Supplemental Figure 1E). These data reinforce previous observations suggesting HP1 proteins regulate a neurodevelopmental transcriptional program (Font-Burgada *et al*. 2008; Ostapcuk *et al*. 2018) and suggest that in addition to their role in formation of chromatin structures, HP1 proteins may also regulate a transcriptional program regulating chromosome organization.

### Cooperative HP1 binding is a better indicator of transcriptional activation than broader chromatin domains

To better understand what factors are associated with HP1-related transcriptional activation, we examined associations between transcriptional status across chromatin states and broader chromatin domains. First, we categorized HP1 targets and non-targets as either heterochromatic or euchromatic to see if differences in transcriptional activity associated with HP1 binding were context specific. Higher expression levels at HP1 target genes was found to be consistent across both heterochromatin and euchromatin (Figure 2A-F). All three HP1 family members were associated with higher expression levels at target genes within euchromatin (Figure 2A-C, p < 2.2e-16, Mann-Whitney). This relationship appeared to be consistent between heterochromatic target and non-target genes (Figure 2D-F). However, the small number of non-target HP1 genes within heterochromatin prevents statistical evaluation of this observation. This finding suggests that knowledge of surrounding chromatin context does not predict whether HP1 proteins exhibit repressive or activating effects on transcription.

**Figure 2.**
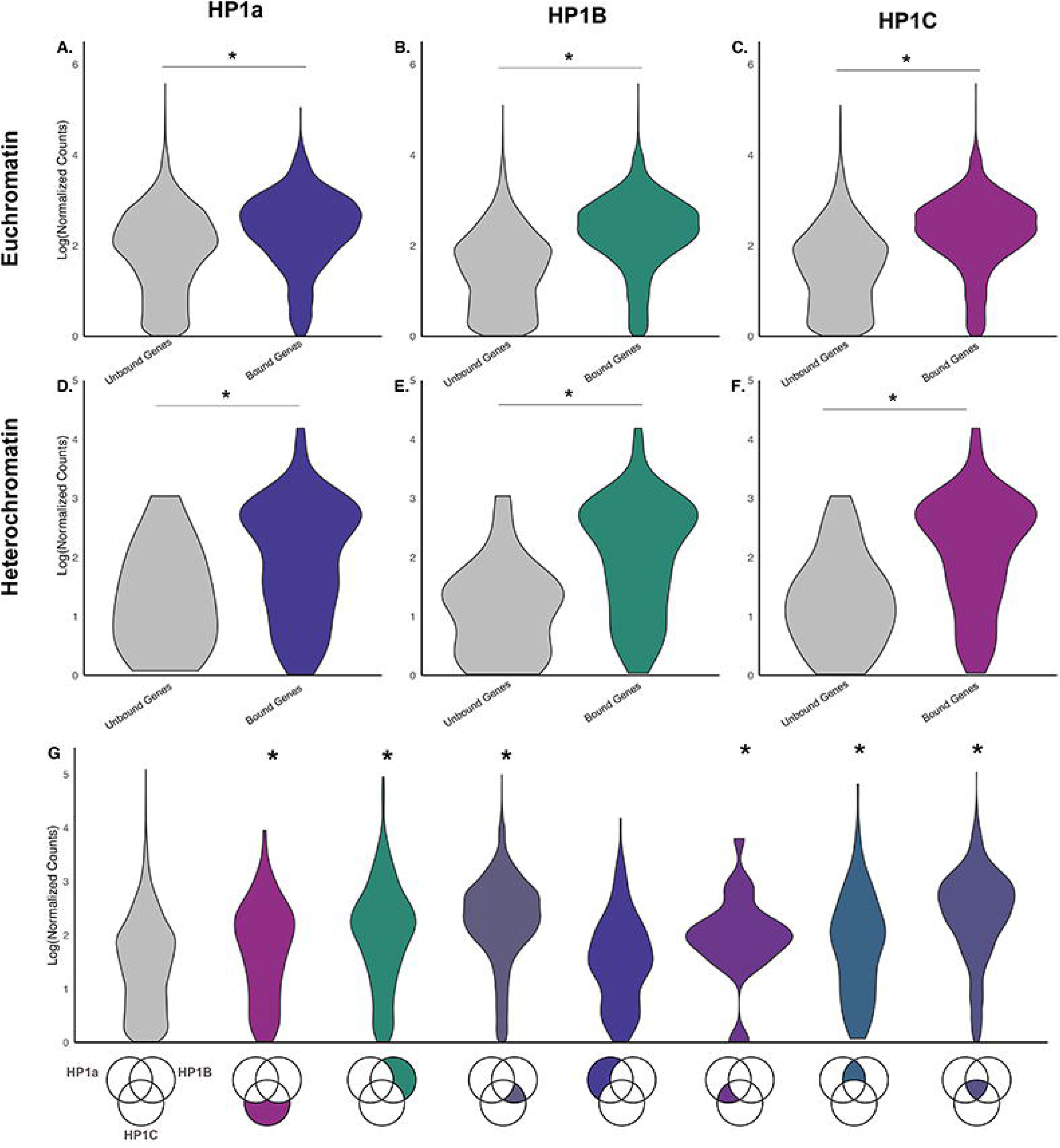
Gene expression analysis of HP1 binding targets across chromatin states and different combinations of HP1 proteins. HP1 binding target (“bound genes,” colored) are expressed significantly higher than non-target genes (“unbound genes,” grey) as measured by Log(1)(TPM) average from two biological replicates (Y-axis, *:p < 2.2e-16, Mann-Whitney). Data from euchromatin for HP1a (A), HP1B (B), and HP1C (C), as well as heterochromatin with HP1a shown in (D), HP1B in (E), and HP1C in (F). (G) Expression level comparison for genes bound by different combinations of HP1 proteins, with HP1 combination indicated in the Venn diagram below the violin plots. Genes bound by a combination of HP1a, HP1B and HP1C or by a combination of HP1B and HP1C are expressed significantly higher than HP1 non-target genes (*:p < 2.2e-16, Mann-Whitney), while genes bound exclusively by HP1a do not exhibit higher expression (p = .226,, Mann-Whitney).

Next, we examined whether considering combinations of HP1 family members present at gene promoters may better predict transcriptional activity. We found that genes bound by particular combinations of HP1 proteins exhibited highest levels of transcriptional activation. Genes bound by either all three HP1 proteins or genes bound by HP1B and HP1C, but not HP1a, exhibited the highest levels of expression compared to genes unoccupied by any HP1 proteins (Figure 2G, p < 2.2e-16, Mann-Whitney). However, genes bound exclusively by HP1a did not exhibit a significant difference in expression compared to genes unbound by any HP1 protein (p = .26, Mann-Whitney). This finding suggests increased expression observed at HP1a binding targets (Supplementary Figure 1A) is driven by genes bound by combinations of HP1 proteins. In summary, the combination of HP1 proteins present at a gene’s promoter is a better predictor of transcriptional activity than knowledge of the surrounding chromatin context.

### HP1 target promoters are enriched for DNA sequence motifs

HP1a binding to gene promoters has been suggested to be independent of its H3K9me2/3 reader activity (Cryderman *et al*. 2005), and HP1C is known to be targeted to chromatin by the DNA binding Zinc Finger transcription factors WOC and ROW (Font-Burgada *et al*. 2008; Kessler *et al*. 2015; Di Mauro *et al*. 2020). Therefore, we performed a motif analysis(Bailey *et al*. 2015) of promoters of HP1 binding targets to identify putative regulatory sequences that may be important for targeting HP1 to protein-coding genes. We looked for enriched motifs in HP1-bound promoters controlling against unoccupied promoters, defining the promoter as the region 250 bp upstream of the TSS. We limited our analysis to the top five enriched motifs in each promoter set. We identified a common, enriched motif present in each set of promoters occupied by HP1a, HP1B, and HP1C (motif ATCGATA, Supplemental Figures 1G-I). This motif was identified previously as a housekeeping core promoter element known as the DNA Replication-related Element (DRE). In *Drosophila*, DRE is known to be recognized by DNA Replication-related Element Factor, the Non-Specific Lethal Complex, and some members of the basal transcription machinery as a key step in transcription initiation (Hochheimer *et al*. 2002; Ohler *et al*. 2002; Juven-Gershon and Kadonaga 2010; Feller *et al*. 2012). The identification of core regulatory elements in promoters of HP1-bound genes raises the possibility that HP1 proteins may mediate transcriptional activation through interactions with core transcriptional machinery.

### Genomic distributions of HP1 proteins in S2 cells match larval distributions

To determine if the results from the analysis of the HP1 binding landscape in third instar larvae is representative of other cell types, we also analyzed available HP1a, HP1B and HP1C binding profiles from Drosophila S2 cells (Ho *et al*. 2014) (Supplemental Figure 2). We measured the degree of colocalization between HP1 proteins in S2 cells and found that 72% of genes bound by HP1a were also bound by HP1B and HP1C, while 17% of HP1a binding targets were bound exclusively by HP1a (Supplemental Figure 2A), recapitulating observations form larvae which showed a large number of genes bound by all three HP1 proteins. HP1B shared 58% of its binding targets with HP1C but not HP1a, as opposed to 18% of HP1B binding targets being shared with both HP1C and HP1a. These results demonstrated that the high degree of shared binding sites between HP1B and HP1C was present in both S2 cells and larvae. 21% of HP1B binding targets were bound exclusively by HP1B (Supplemental Figure 2A). HP1C shared 65% of its genic binding targets with HP1B but not HP1a as opposed to 20% of HP1C binding targets shared by all three proteins (Supplemental Figure 2A). 12% of HP1C targets were exclusively bound by HP1C. The magnitude of these relationships matches observations from the larval datasets. Similar to the binding data from third instar larvae, data from S2 cells demonstrate that a majority of HP1 family binding targets are bound by a combination of different HP1 proteins.

Next, we examined whether HP1 proteins also could be differentiated by their tendency to localize to different functional DNA sequence elements in S2 cells. We again examined five types of annotated sequence elements analyzed earlier, and the results from S2 cells mirror our findings from larvae. HP1a bound 53% of all annotated TEs and 78% of all annotated repeats, while HP1B and HP1C bound less than 1% of all annotated TEs and repeats (Supplemental Figure 2B). HP1B and HP1C were observed at OriCs more often than HP1a (47% and 45% versus 24%; Supplemental Figure 2B). HP1B and HP1C also were associated more frequently with genes than HP1a; HP1a bound 16% of all genes, while HP1B and HP1C bound 26% and 23% of all genes, respectively (Supplemental Figure 2B). A similar trend was observed at enhancers, where HP1a occupied 8% of all annotated enhancers while HP1B and HP1C bound 24% and 26% of all annotated enhancers, respectively. Overall, associations between HP1 proteins and DNA sequence elements in S2 cells matched the findings from third instar larvae and help to differentiate functions among Drosophila HP1 family members.

We next examined the broad chromatin context of genic HP1 binding targets in S2 cells. We found that HP1a was significantly enriched at targets within heterochromatin as well as on chromosome four, binding almost 100% of all heterochromatic genes and approximately 75% of genes located on chromosome four, while only binding approximately 12% of euchromatic genes (Supplemental Figure 2C, p < 2.2e-16, chi square test). HP1B and HP1C also were enriched significantly within heterochromatin and on chromosome four, but to a lesser extent. HP1B bound approximately 50% and 28% of all heterochromatic and chromosome four genes while only binding 25% of all euchromatic genes (Supplemental Figure 2D, p = 1.078e-08, chi square test). HP1C bound 50% and 28% of all heterochromatic and chromosome four genes while only binding 25% of all euchromatic genes (Supplemental Figure 2E, p = 3.022e-14, chi square test). These data again highlight the tendency of HP1 proteins to localize to heterochromatic regions of the genome but also demonstrate that they have binding targets throughout both chromatin compartments. When comparing enriched domain width across compartments, we did detect significantly larger HP1a binding regions within heterochromatin and on chromosome four than within euchromatin (Supplemental Figure 2F, p < 2.2e-16, Mann Whitney), but did not detect a difference for peak size across chromatin contexts for HP1B or HP1C peaks (Supplemental Figures 2G-H). These data again highlight the tendency of HP1 proteins to localize to heterochromatic regions of the genome but also demonstrate that they have binding targets throughout both chromatin compartments. Together, the analysis of HP1 binding data from S2 cells suggests that patterns of binding are similar in chromatin from different sources.

### HP1 genic targets reside in particular chromatin states

Chromatin frequently is classified into higher-order states beyond heterochromatin and euchromatin based on the varying compositions of histone modifications and chromatin-binding proteins (Filion *et al*. 2010; Kharchenko *et al*. 2011). To gain a better understanding of the localization patterns of the different HP1 family members, we determined the extent to which they targeted genes in nine different chromatin states in Drosophila S2 cells defined by the modENCODE consortium (Kharchenko *et al*. 2011). In general, we found that a majority of protein-coding genes reside in chromatin states one, two, three, four, and nine, which correspond to the euchromatic compartment of the genome (Supplemental Figure 2I). We identified enrichment of all HP1 family members at protein coding genes in heterochromatic states six, seven, and eight. Of the euchromatic states, all three HP1 proteins were enriched in state five, which is marked by significant enrichment of H4K16ac and is distributed prominently throughout the Drosophila X chromosome. Meanwhile, we found that all three HP1 proteins were depleted within state nine, which is depleted for most chromatin modifications and proteins and comprises 40% of the Drosophila genome. Depletion of HP1 proteins in this state is consistent with their function as epigenetic readers. We also observed depletion of HP1 targets in euchromatic states one, two and three. These results further strengthen the association of the HP1 family with transcriptionally active chromatin domains.

### HP1 genic targets are enriched for active histone modifications regardless of broader chromatin context

Given that HP1 binding targets are transcriptionally active and that HP1 proteins localize to targets within chromatin states that are both permissive and restrictive to transcription, we sought to profile chromatin marks at HP1 binding targets at increased resolution. A defining structural feature of the HP1 family is the presence of a chromo-domain which permits recognition and binding of H3K9me2/3 (Eissenberg and Elgin 2014). However, the genomic distributions of HP1a and other HP1 proteins are not strictly defined by H3K9me2/3 recognition as HP1 proteins display H3K9me2/3 independent localization (Greil *et al*. 2003; Figueiredo *et al*. 2012). Therefore, we characterized histone methylation patterns at promoters of HP1 binding targets and non-targets. We compared the co-localization of different combinations of HP1 proteins with the repressive histone modifications H3K9me2/3 as well as the active histone modifications H3K4me1/3 across different chromatin contexts (Figure 3A-C). In euchromatin, we observed that localization of HP1B and HP1C is largely independent of H3K9me2/3 except in the presence of HP1a (Figure 3A). Euchromatic genes bound exclusively by HP1B or HP1C were depleted for H3K9me2 (p = 1, 1 respectively, hypergeometric test) and H3K9me3 (p = .99, .99 respectively, hypergeometric test). Euchromatic genes bound by HP1B and HP1C were depleted also for both marks (p = 1, 1, hypergeometric test). In contrast, euchromatic genes bound by both HP1a and HP1B were enriched for both H3K9me2 and H3K9me3. (p = 0, 0 hypergeometric test). We also detected significant association between genes bound by both HP1a and HP1C and these marks (p = 0, 6.79e-01, hypergeometric test). Finally, we detected significant H3K9me2/3 enrichment at euchromatic genes bound by all three HP1 proteins as well as genes bound by HP1a exclusively (hypergeometric test). Thus, associations between the HP1 family and H3K9me2/3 at protein-coding genes are detected only in gene sets where HP1a is present.

**Figure 3.**
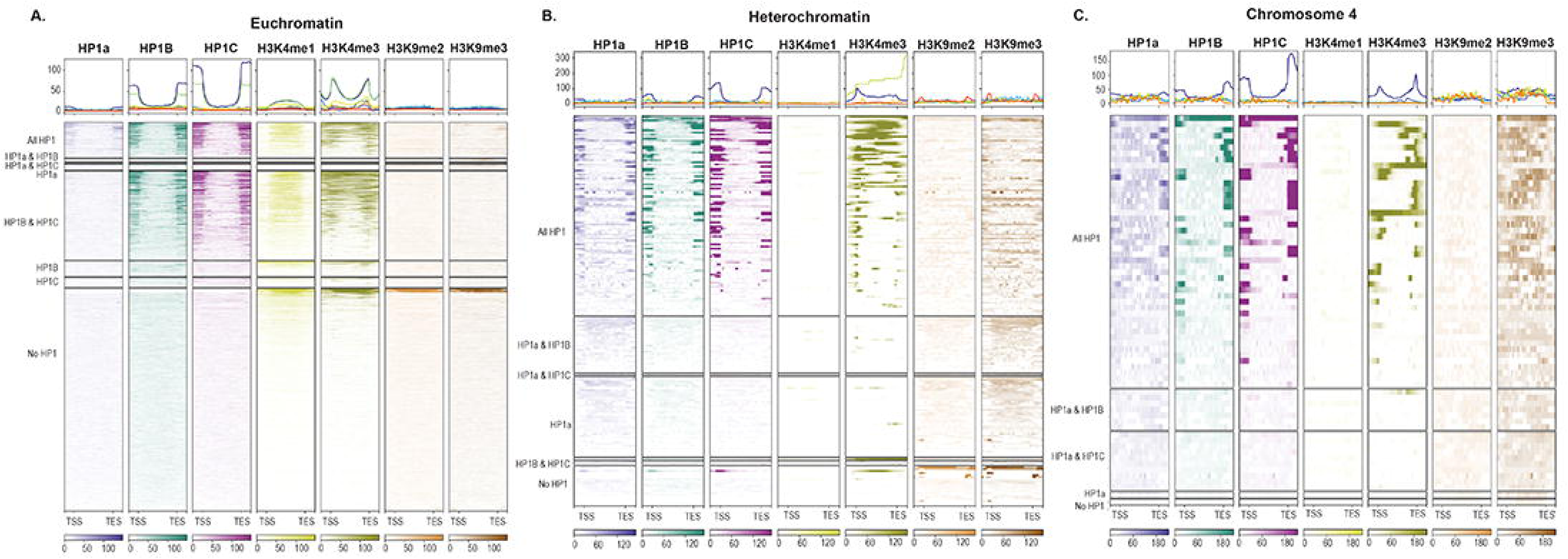
The histone modification context of HP1 target genes across chromatin environments. Average metagene profiles and heatmaps for HP1a, HP1B, HP1C, H3K4me1, H3K4me3,, H3K9me2 and H3K9me3 at HP1 target genes. Genes are classified by the combination of HP1 proteins present at the promoter (left). Only genes that were at least 250 bp away from their nearest neighbor were included for plotting purposes. Color intensity within heatmap reflects coverage at particular loci. Profiles of certain binding combinations of HP1 proteins are excluded due to the small number of genes within that group. (A) Genes within euchromatin. (B) Genes within heterochromatin. (C) Genes located on the 1.2 Mb arm of chromosome four, which is enriched for HP1 proteins.

Because HP1C can be a transcriptional activator in some contexts and HP1 proteins have been associated previously with induced gene expression, we also examined the association between HP1 proteins and active chromatin marks within euchromatin. Interestingly, we detected significant enrichment of histone modifications correlated with active transcription, such as H3K4me1/3 at many of these euchromatic gene groups in addition to H3K9me2/3 enrichment. We detected significant H3K4me1 enrichment at euchromatic genes bound by all three HP1 proteins as well as genes bound by both HP1B and HP1C and genes bound exclusively by HP1C (p = 0, 0, 6.19e-07 respectively, hypergeometric test). We detected H3K4me3 enrichment at euchromatic genes bound by all three proteins, both HP1B and HP1C, or exclusively HP1B (p = 0, 0, 7e-05 respectively hypergeometric test). Overall, analyses of histone modification ChIP-Seq data at euchromatic HP1 family gene targets reinforces a strong association between HP1a and repressive histone modifications while also demonstrating independent localization characterized by active histone modifications.

In addition to studying associations between HP1 binding targets and different chromatin modifications within euchromatin, we also measured associations between HP1 binding targets and different histone methylation marks in heterochromatic contexts. We detected joint enrichment of active and repressive histone modifications at heterochromatic HP1 binding targets (Figure 3B). Heterochromatic genes bound by all three HP1 proteins were enriched for all histone modifications analyzed (p = 0, 0, 0, 0 hypergeometric test). We also detected significant enrichment of H3K4me1 at heterochromatic genes bound by both HP1B and HP1C (p = 0 hypergeometric test). Due to a small number of genes, we were unable to assess enrichment for other combinations of HP1 proteins within heterochromatin. However, enrichment of H3K4 methylation at heterochromatic HP1 targets is in agreement with previous data demonstrating that heterochromatic HP1a binding targets are actively transcribed.

Finally, we examined association patterns between HP1 proteins and histone modifications on chromosome four. We again detected significant enrichment of all histone modifications analyzed at chromosome four genes bound by all three HP1 proteins (p = 0, 0, 0, 0, hypergeometric test). In addition, we detected enrichment of H3K4me3 at genes bound by HP1B and HP1C as well as genes bound by HP1C exclusively (p = 0, 0, hypergeometric test). Limited sample size prevented enrichment analysis of other HP1 protein combinations. Histone modification data from chromosome four again reinforces the association between HP1 proteins and active chromatin states.

### HP1 binding targets display signatures of promoter proximal RNA polymerase pausing

All three somatic Drosophila HP1 proteins co-immunoprecipitate with both subunits of the Facilitates Chromatin Transcription (FACT) complex, which promotes transcriptional elongation (Kwon *et al*. 2010). Furthermore, HP1C has been implicated previously in release from promoter proximal pausing (Kessler *et al*. 2015), and HP1a and HP1C have been associated with pausing at transcribed genes (Sakoparnig *et al*. 2012). However, the extent to which this relationship depends on the cooperative activity of other HP1 proteins has not been examined. To better understand the association between the HP1 protein family and transcriptional pausing by RNA polymerase II (referred to as ‘pausing’), we compared RNA polymerase II (RPII) dynamics at HP1 target and non-target genes using available next generation sequencing datasets. Metagene profiles of RPII ChIP-Seq data demonstrated that HP1 target genes generally displayed a higher 5’ RPII signal peak in addition to overall increased RPII recruitment. (Figures 4A-C). To quantify this relationship, we calculated pausing indices (Muse *et al*. 2007; Larschan *et al*. 2011). Here, each gene is divided into two regions (Figure 4D). A pausing index can be calculated by dividing the read density in the 5’ region over the read density in the mid-gene region. Pausing indices allow for the evaluation of RPII dynamics using next-generation sequencing datasets. We calculated pausing indices for HP1 target and non-target genes using available RPII ChIP-Seq data from Drosophila third instar larvae. HP1a, HP1B, and HP1C target genes all had significantly higher pausing indices than non-target genes (Figure 4E-G, p < 2.2e-16; Mann-Whitney). This finding validates previously observed associations between HP1a and HP1C with promoter proximal pausing and is the first evidence of a possible role for HP1B in promoter proximal pausing. Next, we examined RPII dynamics at protein coding genes with different combinations of HP1 proteins present at the promoter. We overlaid RPII ChIP-Seq metagene profiles of each of these gene groups (Figure 4H). This visualization revealed that the group of genes bound exclusively by HP1a (blue line) did not contain a higher 5’ peak compared to genes without HP1 proteins present at the promoter (gray line). Genes bound exclusively by HP1C (bright purple) or by a combination of HP1a and HP1C (dark purple) displayed moderately increased RPII recruitment at the promoter compared to genes unoccupied by HP1 family members. Genes bound by a combination of HP1 family members that includes HP1B had highest RPII 5’ peaks and increased RPII recruitment over the gene body compared to other gene groups. Analysis of pausing indices of these gene groups further highlighted these differences. Pairwise Mann-Whitney U tests with false discovery rate (FDR) corrections demonstrated that most HP1 family combinations were distinct (Figure 4I). Average pausing indices were highest in gene groups occupied by an HP1 combination that included HP1B. Of these groups, genes bound by all three HP1 proteins had the highest overall pausing indices. Meanwhile, genes bound exclusively by HP1a had the lowest pausing indices of any combination of HP1 proteins and were not significantly different from genes without HP1 family members at the promoter. Genes bound by HP1C exclusively also had lower pausing indices than other HP1 family combinations but were significantly different from genes without any HP1 proteins. Notably, the group of genes bound by HP1a and HP1C is difficult to interpret due to the small sample size of this group compared to other gene groups (n = 20). Overall, these data present strong evidence for the importance of cooperative activity among HP1 family members, particularly HP1B, in observed associations between the HP1 family, promoter proximal pausing and transcriptional activation.

**Figure 4.**
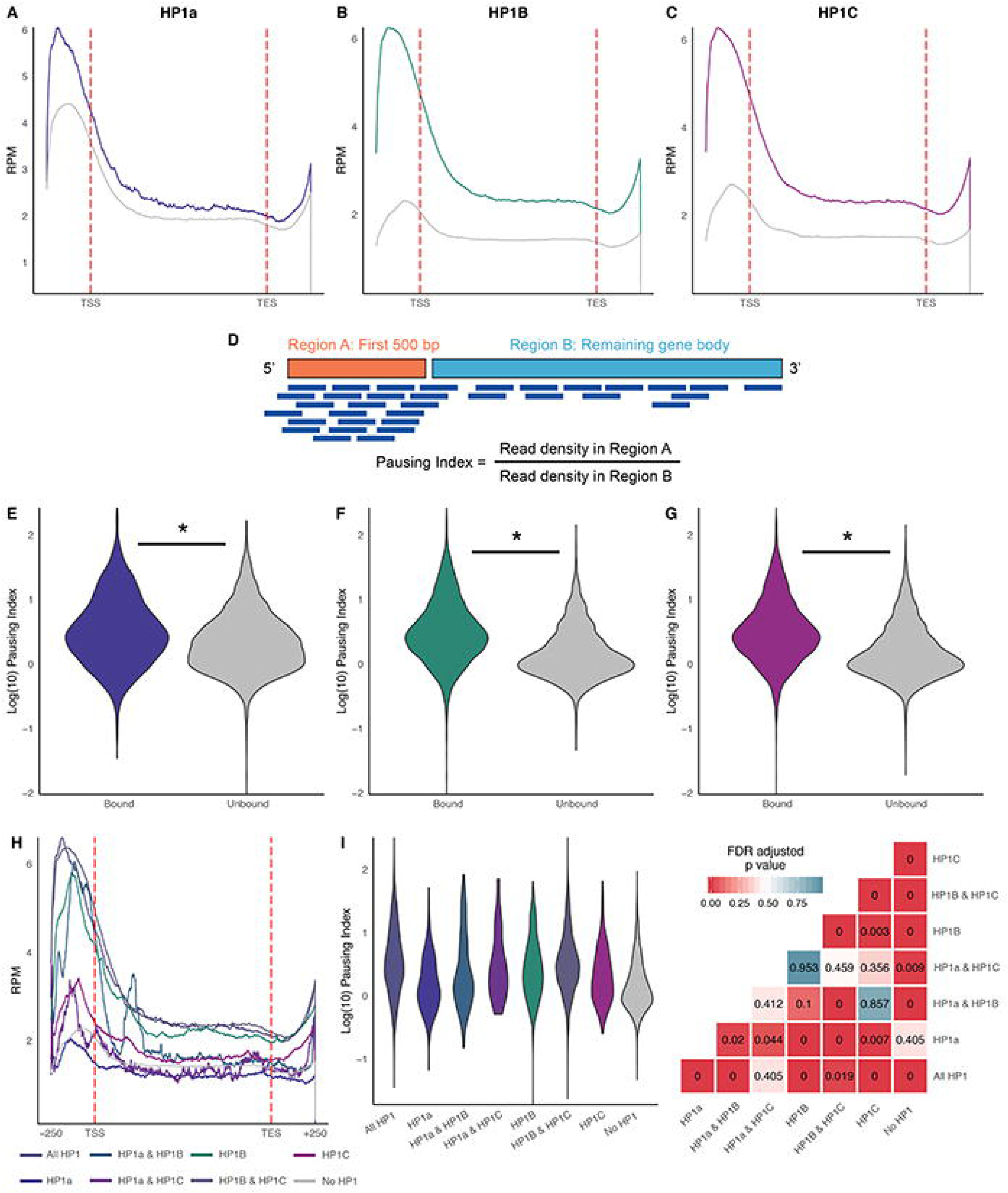
Increased promoter proximal pausing at HP1 binding targets. (A-C) RPII ChIP-Seq metagene profiles from third instar larvae demonstrate that HP1a, HP1B, and HP1C binding targets have higher 5’ peaks in RPII ChIP-Seq signal as well as increased RPII recruitment over the gene body. X-axis: scaled position from TSS to TES; Y-axis: RPM. (D) Illustration of pausing index calculation used to quantify RPII dynamics across HP1 binding states. (E-G) HP1 binding targets (“Bound”) had significantly higher pausing indices compared to genes that were not occupied by HP1 at the promoter (“Unbound”). Y-axis Log_10_(Pausing Index). (E) – HP1a. (F) – HP1B, (G) – HP1C. (H) Metagene profiles of genes grouped by different combinations of HP1 proteins. X axis: scaled position from TSS to TES; Y-axis: RPM. (I) Violin plot of pairwise comparisons of different HP1 proteins, FDR adjusted p values for pairwise comparisons are presented in corresponding heatmap. X-axis: Combination of HP1 proteins. Y-axis: Log_10_(Pausing Index).

To validate these findings, we sought whether relationships between HP1 family members and pausing were observable through an analysis of the nascent transcriptome. We analyzed available Global Nuclear Run-On followed by next generation sequencing (GRO-Seq) data from Drosophila S2 cells as an orthogonal approach to RPII ChIP-Seq to assess RPII dynamics at HP1 family target genes (Supplemental Figure 3). We generated GRO-Seq metagene profiles of HP1 target and non-target genes in S2 cell culture from available data (Larschan *et al*. 2011) (Supplemental Figure 3A-C). Metagene profiles recapitulated observations from RPII ChIP-Seq data. HP1a, HP1B and HP1C target genes had higher 5’ peaks and increased signal over the gene body than non-target genes. We again calculated pausing indices to quantify these observations. Pausing index analysis again recapitulated HP1a, HP1B and HP1C target genes all had higher pausing indices than respective non-target genes (Supplemental Figures 3D-F). These results demonstrate that the association between HP1 family members and active transcription at TSSs of protein-coding genes can be observed not only through measurements of RPII positioning but also through analysis of the nascent transcriptome. To follow up this analysis, we examined how nascent transcription signatures vary across genes bound by different combinations of HP1 proteins. We analyzed pausing indices across these different gene groups using pairwise Wilcoxon tests with FDR correction (Supplemental Figure 3H). This analysis produced similar results to our analysis of RPII ChIP-Seq data (Figure 4I). Genes bound by all three HP1 proteins or by a combination of HP1B and HP1C again had the highest mean pausing indices. Genes bound by other combinations of multiple HP1 proteins – HP1a and HP1B, HP1a and HP1C – as well as genes bound exclusively by HP1C comprised a middle tier of pausing indices. Genes bound exclusively by HP1C or HP1a or genes not occupied by HP1 family members at their promoter had the lowest pausing indices (Figure 4I). These observations are well illustrated by metagene profiles of GRO-Seq data across these gene groups (Figure 4H). Overall, our analysis of nascent transcription dynamics at HP1 target genes validates key findings of our analysis of RPII ChIP-Seq data. Namely, that particular combinations of HP1 proteins are consistently and strongly associated with transcriptional activation.

### Associations between HP1 and pausing are consistent across heterochromatin and euchromatin

To better understand the association between the HP1 family and transcriptional pausing, we next set out to investigate whether surrounding chromatin context differentiated between the degree of pausing at HP1 binding targets. We generated metagene profiles and compared average pausing indices for HP1a, HP1B, and HP1C binding targets in both heterochromatic and euchromatic contexts from RPII ChIP-Seq data from third instar larvae (Supplemental Figure 4). Metagene profiles of HP1 target and non-target genes in both contexts demonstrated that higher RPII 5’ peaks and increased RPII recruitment was consistent at HP1 binding targets across chromatin contexts (Supplemental Figures 4A-B, E-F, I-J). Quantifying this observation with pausing indices found that binding targets of all three HP1 proteins exhibited significantly higher pausing indices than unbound genes in euchromatic contexts (Supplemental Figures 4D, H and L, p < 2.2e-16, Mann-Whitney). We also detected significantly increased pausing indices in heterochromatin at HP1C target genes (Supplemental Figure 4K, p = 0.002, Mann-Whitney) and at HP1B target genes (Supplemental Figure 4G, p = 0.002, Mann-Whitney). We did not detect significant differences in pausing indices between HP1a target and non-target genes in heterochromatin; however, this result may be due to the small (n = 12) sample size of non-target genes in this chromatin context (Supplemental Figure 4C). Overall, our analysis of RPII ChIP-Seq dynamics across chromatin contexts suggests that HP1 binding activity at TSSs is associated with a consistent functional outcome of increased pausing and gene expression in both heterochromatin and euchromatin.

To validate these findings, we again analyzed orthogonal GRO-Seq data to see whether the association between HP1 binding and increased pausing was visible through analysis of the nascent transcriptome. We generated GRO-Seq metagene profiles for HP1a, HP1B and HP1C binding targets in heterochromatin and euchromatin. All three HP1 family members had increased signal at the 5’ gene end as well as over the gene body in both contexts (Supplemental Figure 5A-B, E-F, I-J). Quantifying metagene profiles with pausing indices recapitulated findings from RPII ChIP-Seq data. HP1a binding targets were paused significantly compared to non-targets in euchromatin, but not heterochromatin. HP1B and HP1C binding targets were paused significantly compared to non-targets in both heterochromatin and euchromatin. These results again indicate that the broader surrounding chromatin context is not predictive of transcriptional activity.

### HP1-bound genes are enriched for histone modifications associated with enhanced transcriptional elongation

In addition to measuring pausing indices at HP1 binding targets, we also investigated whether HP1-bound genes were enriched for particular histone modifications correlated with RPII elongation (Veloso *et al*. 2014; Chen *et al*. 2018). We measured the associations between HP1 family member binding with three histone modifications: Histone 4 lysine 20 monomethylation (H4K20me1), Histone 2B lysine 120 ubiquitination (H2B-ubi) and histone 3 lysine 79 monomethylation (H3K79me1) using available ChIP-Seq data (Supplemental Figure 6). We evaluated enrichment patterns across different chromatin contexts to observe whether associations were specific to a particular broader chromatin context. Overall, patterns of association were very similar for all three histone modifications. In euchromatin, we detected significant enrichment of H2B-ubi at genes bound by all three HP1 proteins as well as genes bound by HP1B and HP1C (p = 0, 0, respectively, hypergeometric test). We found this same enrichment pattern with respect to H4K20me1, in addition to significant enrichment of H4K20me1 at genes bound exclusively by HP1B (p = 0, 0, 0.012, hypergeometric test). We found significant enrichment of H3K79me1 at euchromatic genes bound by all three HP1 proteins, genes bound by HP1a and HP1C, genes bound by HP1B and HP1C as well as genes bound exclusively by HP1B (p = 0, 0, 0, 5.34e-13, hypergeometric test). Meanwhile in heterochromatin, we detected significant enrichment for H2B-ubi at genes bound by all three HP1 proteins as well as genes bound exclusively by HP1C (p = 0, 0, respectively, hypergeometric test). We also detected significant H4K20me1 enrichment at genes bound by all three HP1 proteins or genes bound exclusively by HP1C within heterochromatin (p = 0, 0, hypergeometric test). With respect to H3K79me1, we detected significant enrichment at genes bound by all three HP1 proteins, genes bound by HP1B and HP1C, and genes bound by HP1C exclusively (p = 0, 0.013, 0, hypergeometric test). Finally, on chromosome four, we detected significant H2B-ubi enrichment at genes bound by all three HP1 proteins, genes bound by HP1B and HP1C, and genes bound exclusively by HP1C (p = 0, 0, 0, hypergeometric test). We detected this same enrichment pattern across gene groups with respect to H4K20me1 (p = 0, 0, 0, hypergeometric test) as well as H3K79me1 (p = 0, 0, 0, hypergeometric test). Observed enrichment of these histone modifications at HP1 target genes supports our findings that the colocalization of HP1 proteins is strongly associated with RPII activity and increased gene expression.

### HP1 depletion impacts gene expression

To understand how HP1 proteins regulate gene expression, we integrated three RNA-Seq datasets of HP1 knockout mutants. We utilized available datasets of *Su(var)205* and *HP1b* knockout mutants (Riddle *et al*. 2012; Mills *et al*. 2018) and generated a novel library to study gene expression in an *HP1c* knockout mutant (Supplemental Figure 7). We then compared differentially expressed genes across all three datasets to better understand the set of genes regulated by the HP1 family. We found that depletion of HP1a and HP1B resulted in upregulation of a large number of genes and a smaller quantity of downregulated genes, while depletion of HP1C resulted in both up- and downregulated gene expression at approximately equal levels (Supplemental Figure 7). Next, we examined changes in gene expression upon HP1 depletion at genes bound by HP1 proteins. We found that 48.83% of HP1a bound genes were differentially expressed upon HP1 depletion. A majority of expression changes observed upon HP1a depletion appear to be due to secondary effects, evidenced by the fact that only 19.73% of differentially expressed genes were binding targets (Supplemental Figure 7G). In contrast, we found that HP1B and HP1C binding targets constituted a small majority of differentially expressed genes, although only a small percentage of binding targets was differentially expressed (Supplemental Figures 7H-I). Upon HP1B depletion, 50.95% of differentially expressed genes are bound by HP1B under wildtype conditions, although only 17.50% of binding targets were differentially expressed (Supplemental Figure 7H). Similarly, 52.78% of differentially expressed genes upon HP1C depletion are genes bound by HP1C under wildtype conditions, but only 16.57% of HP1C binding targets are differentially expressed upon HP1C depletion (Supplemental Figure 7I). Therefore, while a majority of HP1B and HP1C binding targets do not experience significant changes in expression upon depletion of either respective protein, those genes which are differentially expressed constitute a small majority of observed transcriptional changes.

### Depletion of individual HP1 proteins reveals roles for HP1 family members in promoter proximal pausing

To better understand the impact of HP1 binding on promoter proximal pausing, we measured pausing indices in knockout mutants for HP1a, HP1B, and HP1C using RPII ChIP-chip data from third instar larvae made available by the modENCODE consortium (Ho *et al*. 2014). We used an alternative pausing index calculation that is compatible with ChIP-chip datasets (Zeitlinger *et al*. 2007) and again analyzed the pausing indices at all genes bound by each HP1 protein as well as pausing indices bound by different combinations of HP1 proteins. Using this modified calculation, we were able to detect significantly increased pausing at HP1a, HP1B and, HP1C target genes in wild-type Drosophila third instar larvae (Figures 5A, C, and E). Overall, significantly increased promoter proximal pausing at HP1 target genes was maintained in respective knockout mutants (Figures 5B, D and F). This observation is consistent with a model where HP1 proteins cooperate to regulate transcription and exhibit a degree of functional redundancy at TSSs.

**Figure 5.**
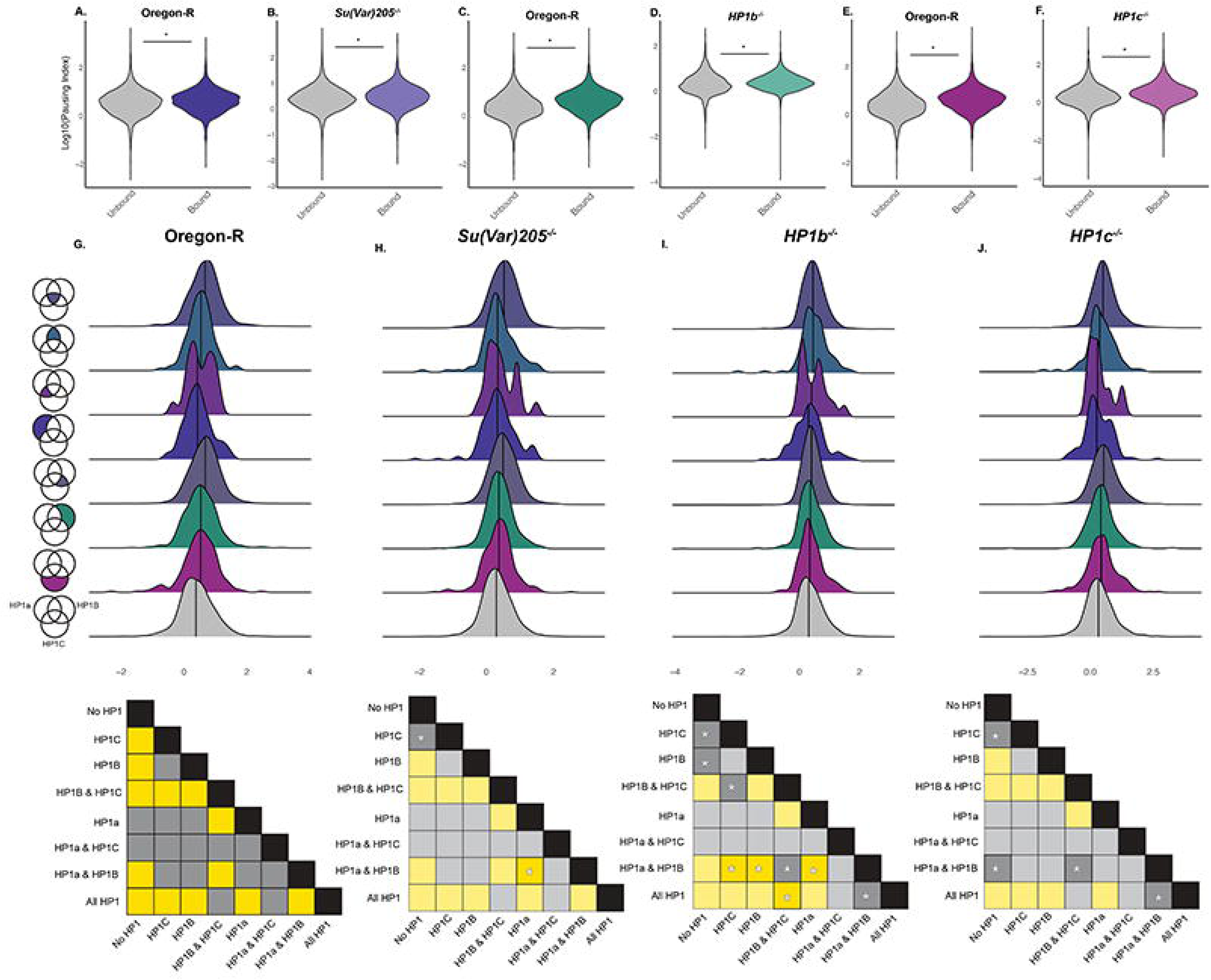
Depletion of HP1 proteins alters dynamics of promoter proximal pausing. (A-E) HP1 binding targets (colored) had significantly higher pausing indices than non-targets (grey) as measured by RPII ChIP-chip data in wild type (darker colored plots, left side) and HP1 null mutant (lighter colored plots, right side) third instar larvae. (A – B) HP1a targets in wild type (p = 1.05e-10, Mann-Whitney) and null mutant (p < 2.2e-16, Mann-Whitney) larvae (Blue). (C – D) HP1B targets in wild type (p < 2.2e-16, Mann-Whitney) and null mutant (p < 2.2e-16, Mann-Whitney) larvae (Green). (E – F) HP1C targets in wild type (p < 2.2e-16, Mann-Whitney) and null mutant larvae (p < 2.2e-16, Mann-Whitney) (Purple). X-axis: Binding classification. Y-axis: Log_10_(Pausing Index). (G-J) Ridge plot comparing distributions of pausing indices when genes are grouped by the combination of HP1 proteins present at the TSS. X-axis: Log_10_(Pausing Index). Y-axis: Frequency. Genes are grouped by the combination of HP1 proteins present at the TSS, as denoted by Venn diagrams. Results of pairwise comparisons between groups are summarized in grid below plots. Significant differences between groups are highlighted yellow (FDR adjusted cutoff p < 0.05). Opaque squares denoted with * in (H-J) signify pairwise comparisons observed to deviate from results in wild type larvae. (G) – wild type. (H) – *Su(Var)205* null mutant. (I) – *HP1b* null mutant. (J) – *HP1c* null mutant.

Analysis of pausing indices across genotypes suggests binding of HP1B and HP1C may be particularly important for transcriptional regulation by HP1 family members. To gain insight into individual functions of HP1 proteins in transcriptional regulation, we decided to examine how pausing indices changed across HP1 null mutants at genes bound by different combinations of HP1 proteins (Figures 5G-J). We first compared pausing indices across HP1 binding groups in the wildtype dataset with functional copies of all three somatic HP1 genes to better appreciate how the groups relate to each other in the ‘wild type’ condition. A Kruskal-Wallis test confirmed that there were significant differences in pausing indices across HP1 binding groups (X^2^ = 526.62, p < 2.2e-16) which we followed up with pairwise Wilcoxon tests with FDR correction to examine pairwise differences. We found a total of 13 significantly different pairwise comparisons between different HP1 binding groups which roughly partitioned the groups into three tiers (Figure 5G). Genes that were not bound by any HP1 proteins did not have a significantly different pausing index compared to genes bound exclusively by HP1a and these groups had the lowest average pausing indices. A middle tier of groups was comprised of genes bound exclusively by HP1C, genes bound exclusively by HP1B, and genes bound by a combination of HP1a and HP1B but lacking HP1C. Groups in this tier had intermediate average pausing index values. Finally, genes bound by both HP1B and HP1C as well as genes bound by HP1a, HP1B, and HP1C did not exhibit significant differences in their pausing indices, and these genes had the highest average pausing indices. (The group of genes bound by HP1a and HP1C were not compared in pairwise comparisons because the bimodal distribution of pausing indices in this group precludes necessary assumptions for statistical inference). These results reinforce prior data suggesting that the colocalization of HP1B and HP1C may be particularly important for the increased pausing and increased expression that has been previously associated with HP1 binding.

Depletion of HP1a results in minor impacts to pausing indices at HP1 target genes. We repeated the above analysis in HP1a null larvae to infer the importance of HP1a in transcriptional regulation (Figure 5H). A Kruskal-Wallis test established significant differences in pausing indices across groups of genes bound by different combinations of HP1 family members (X^2^ = 564.96, p < 2.2e-16). Follow-up of pairwise comparisons using Wilcoxon tests with FDR correction revealed two pairwise comparisons that deviated from the wildtype genotype. Genes bound exclusively by HP1C no longer exhibited significantly increased pausing indices upon depletion of HP1a. Instead, this group of genes now occupied the lowest tier of pausing indices. The second novel difference was that genes bound by HP1a and HP1B had significantly higher pausing indices than genes bound exclusively by HP1a upon HP1a depletion. However, this change did not meaningfully move this group of genes into a new tier of pausing indices. While depletion of HP1a produced some changes in promoter proximal pausing at genes bound by certain combinations of HP1 family members, overall effects were minimal.

In contrast to HP1a depletion which resulted in minimal effects on promoter proximal pausing, depletion of HP1B disrupted promoter proximal pausing on a larger scale. A Kruskal-Wallis test of pausing indices across groups of genes bound by different combinations of HP1 proteins confirmed significant differences between groups (X^2^ = 137.12, p < 2.2e-16). Pairwise Wilcoxon comparisons with FDR correction identified a total of eight comparisons that differed from their respective result in the wildtype genotype. Genes bound exclusively by HP1B or exclusively by HP1C no longer displayed significantly higher pausing indices compared to genes with no HP1 proteins present, contributing to the lowest tier of gene groups ranked by pausing-indices. Additionally, genes bound by a combination of HP1B and HP1C were not significantly different from genes bound exclusively by HP1C, although the former were still significantly different from genes with no HP1 proteins at all. Genes bound by HP1B and HP1C no longer occupied the highest tier of pausing indices upon depletion of HP1B and also exhibited significant differences with genes bound by all three HP1 proteins. The relationship between genes bound by HP1a and HP1B exhibited the most change in this genotype compared to pairwise comparisons in wildtype. Upon depletion of HP1B, these genes had higher pausing indices compared to genes bound exclusively by HP1a, HP1B, or HP1C. However, these genes were not significantly different from genes bound by HP1B and HP1C. These data suggest that HP1B may be particularly important for relationships between HP1 family members when regulating transcription start site activity and that HP1 family members may functionally compensate upon HP1B depletion.

Depletion of HP1C minimized differences in pausing indices across groups of HP1 genes. A Kruskal-Wallis test confirmed significant differences in pausing indices across groups of genes bound by different combinations of HP1 proteins upon depletion of HP1C (X^2^ = 271.35, p < 2.2e-16). Pairwise comparisons using Wilcoxon tests with FDR corrections identified four pairwise comparisons whose relationship differed from the wildtype condition. Each of these comparisons represented a transition from a statistically significant difference to a nonsignificant difference following HP1C depletion. First, genes bound by HP1C were no longer significantly different from genes not bound by HP1 family members. The remaining three comparisons all involved the group of genes bound by HP1a and HP1B. This gene group was no longer significantly different from genes bound by all three HP1 proteins, genes bound by HP1B and HP1C, and genes not bound by HP1 proteins. This observation suggests that the presence of HP1C is important for regulating pausing when different combinations of HP1 proteins are present at transcription start sites.

## DISCUSSION

Here, we analyzed high resolution ChIP-Seq maps of all the somatic *Drosophila* HP1 family members, which raises interesting points about the role of these proteins in gene regulation. We find that all three HP1 proteins bind throughout heterochromatin and euchromatin compartments. With regards to binding behavior at protein-coding genes, while all three HP1 proteins are enriched at genes located within heterochromatin, a majority of their binding targets are located within euchromatin. This finding is true even of HP1a, whose localization often is described as restricted to heterochromatin, as well as HP1C, whose localization tends to be described as restricted to euchromatin. In addition to previously reported enrichment of HP1a on chromosome four, we also detect significant enrichment of HP1B and HP1C. Additionally, the three HP1 proteins share a majority of their binding sites. A gene bound by any HP1 protein is most likely also bound by at least one other family member. This relationship was true across heterochromatin and euchromatin and highlights the need to consider what effect interactions between HP1 proteins have on transcription.

A close examination of HP1 genic binding targets suggests that knowledge of the presence of additional HP1 proteins is a better indicator of transcriptional status than knowledge of the broader surrounding chromatin context. HP1-bound genes are expressed at higher levels than unbound genes across chromatin contexts. Genes bound by all three HP1 proteins or by a combination of HP1B and HP1C are consistently expressed at higher levels across all contexts. HP1-bound genes display a strong association with H3K4me3 across all chromatin contexts but share a context-specific association with H3K9me2/3 within heterochromatin. The independence of HP1 binding to euchromatic genes from H3K9me2/3 matches previously observed data. Cooperative binding of multiple HP1 proteins therefore appears to be a stronger indicator of transcriptional activation than chromatin context.

Signatures of promoter proximal pausing at HP1 binding targets give clues to a potential mechanism of gene activation by HP1 proteins. Here, we report that genes bound by HP1 proteins display higher pausing indices compared to unbound genes. This effect is observed across chromatin states. A pausing index is an indirect measurement of RPII activity that reflects a higher density of RPII at the 5’ end of genes. It is not always clear what factors drive this increased density. For instance, genes with increased pausing durations would be expected to have higher pausing indices and lower expression levels. In contrast, genes with shorter pausing durations but increased initiation frequencies could exhibit high pausing indices in addition to high expression levels (Gressel *et al*. 2017). HP1 binding targets are expressed at higher levels than non-target genes and have strong associations with the active histone modification H3K4me3, in the support of the latter model of increased pausing indices. This observation is supported by observations made by others that HP1 binding targets appear to be both paused and highly transcribed (Sakoparnig *et al*. 2012). Increased pausing indices associated with HP1 binding may be due to the relationship between the HP1 family and the FACT complex, which promotes RPII elongation by removing nucleosomal barriers (Orphanides *et al*. 1998; Kwon *et al*. 2010). Alternatively, increased RPII pausing at HP1 target genes may be regulated through HP1-mediated recruitment of additional factors such as dDsk2 (Kessler *et al*. 2015; Di Mauro *et al*. 2020)Additional evidence is necessary to fully understand the contribution of each HP1 family member to transcriptional activation.

Our analysis of RPII dynamics in single knockout HP1 mutants suggests that interactions between HP1 family members are important in the regulation of gene expression. HP1 targets comprise a majority of differentially expressed genes in *HP1b* and *HP1c* null mutants, and a large fraction of HP1a binding targets are differentially expressed in *Su(var)205* mutants. An analysis of RPII activity at these genes in respective HP1 null mutants supports a model where HP1 proteins promote increased gene expression through regulation of RPII activity. This model is further supported by an observed interaction between HP1 family members and the FACT complex and is consistent with observations of *HP1d* activity in the Drosophila genome (Andersen *et al*. 2017). Our analysis builds on these results by providing insights into how HP1 proteins cooperatively regulate RPII activity in Drosophila somatic cells.

While HP1a and HP1C previously have been implicated in transcriptional activation and promoter proximal pausing individually, ours is the first study to consider how coordinated activity between HP1 proteins may impact gene expression. Additionally, ours is the first study to show genome-wide evidence for a role of HP1B in promoter proximal pausing to induce transcription. Previous studies have suggested that surrounding chromatin contexts may predict whether HP1 proteins have an activating or repressive role at TSSs. However, our genome-wide analysis of HP1 binding targets demonstrates that co-localization of HP1 proteins is a better predictor of whether binding targets are transcribed or repressed than knowledge of surrounding chromatin context. Certain combinations of HP1 proteins, particularly the colocalization of HP1B and HP1C, are strongly associated with active transcription throughout heterochromatin and euchromatin, while HP1a binding on its own is not associated with pausing or transcription. Overall, our analysis highlights the need to consider how HP1 family members work together to regulate gene expression.

## METHODS

### ChIP-Seq Analysis

HP1 binding sites from third instar larvae and S2 cells were downloaded from GEO (see accession numbers in supplementary table 1). Peak genomic coordinates were converted from dm3 to dm6 using the UCSC genome liftOver tool (Kent *et al*. 2002) and compared with annotated protein-coding genes in the Drosophila genome (release 6.25 (Thurmond *et al*. 2019)) to classify genes as bound. Chromatin context boundaries to differentiate heterochromatin and euchromatin were obtained from (Riddle *et al*. 2011). Enrichment of bound genes across chromatin contexts was evaluated using a Chi-square test.

To generate genome-wide binding profiles of HP1 proteins and histone modifications, we downloaded raw sequencing data (see accession numbers in supplementary table 1). Reads were aligned to the dm6 reference assembly using the bwa mem algorithm (version 0.7.16a-r1181) (Li and Durbin 2009). Coverage was calculated with samtools version 1.5 (Li *et al*. 2009) and plotted using Circos (Krzywinski *et al*. 2009). Heatmaps of histone modifications were generated using deepTools (version 3.0.2) (Ramirez *et al*. 2016).

To assess the enrichment of histone modifications at HP1 binding sites, histone modification peak data was downloaded from GEO (Supplementary Table 1) and coordinates were uploaded to UCSC genome liftOver to convert to dm6. Overlap between histone modification peaks and HP1-bound genes was evaluated using a custom python script (Made available at location Y). Enrichment was determined from number of overlaps using hypergeometric tests.

### RNA-Seq analysis

For preparation of transcriptomic data from *HP1c* null mutants, 20 mg of frozen third instar larvae were homogenized, and RNA samples were isolated using Trizol. RNA sample integrity was confirmed by formaldehyde agarose gel electrophoresis. RNA samples were prepared for whole transcriptome sequencing by the UAB Heflin Center for Genomic Science Genomics Core lab. 30-40 million RNA-seq reads were collected per sample using the Illumina Sequencing Platform. We analyzed two RNA-seq samples of the *HP1c* null mutant genotype.

To analyze RNA-seq data, we aligned reads to the dm6 reference genome assembly using STAR aligner (Version #2.5.2) (Dobin *et al*. 2013)and determined transcript counts using HTSeq (version #0.6.1) (Anders *et al*. 2015). Differential expression analysis was performed using DESeq2 (Version #1.22.2) (Love *et al*. 2014). Only genes meeting an FDR (false discovery rate) cut-off of 0.05 were used for downstream analyses. Gene ontology analysis was performed using DAVID (version #6.8) (Huang da *et al*. 2009b; Huang da *et al*. 2009a).

### Motif Analysis

We defined promoter regions as the region covering 250 bp upstream of the TSS to the TSS. Motif analysis of promoter sequences was evaluated using MEME (version 5.1.0), (Bailey *et al*. 2015) searching for the top three hits in each dataset.

### Metagene Profiles

Metagene profiles were calculated using a custom R script. To generate metagene profiles of RPII ChIP-Seq and GRO-Seq data, we first filtered genes shorter than 1 kb. Remaining genes along with the regions 250 bp upstream and downstream were scaled into 1.5 kb. Coverage profiles for each gene were calculated individually and then average before averaging into group profiles.

### Pausing indices

Pausing indices were calculated using a custom R script. To calculate pausing indices, genes were divided into regions: the first five hundred base pairs at the 5’-most end of the gene and the remaining gene body. Read densities were calculated from each region. Genes shorter than 1000 bp or genes with less than three reads aligning to either regions were filtered out of the analysis. Mann-Whitney U tests were used for comparisons across groups.

### Data Availability

The authors state that all data necessary for confirming the conclusions presented in the article are represented fully within the article. All data used in this study are publicly available and referenced in *Materials and Methods*. The Supplemental material will be available at FigShare. R code for the calculation of pausing indices and metagene profiles is available on Github: https://github.com/schoelz-j/schoelz_feng_riddle_2020

## Acknowledgements

The authors would like to acknowledge the support of the Riddle lab members for stimulating discussions and comments on the manuscript. Additionally, the authors would like to thank the University of Alabama at Birmingham Heflin Genomics Core for assistance in generation and analysis of RNA-Seq datasets. This material was supported by NSF CAREER(1552586). Additionally, This material was supported in part by the Alabama State funded Graduate Research Scholars Program (GRSP).

## FIGURE CAPTIONS

**Supplemental Figure 1. Functional annotation of HP1-bound genes in third instar larvae.** (A-C) Violin plot comparing average log(TPM) between HP1 binding targets (“Bound”, color) and non-target genes (“Unbound”, grey) from third instar larvae. (* = p< 0.0002, permutation test) (A) – HP1a; (B) – HP1B; (C) – HP1C. (D-F) Gene ontology analysis of HP1 binding targets (D) – HP1a; (E) – HP1B; (F) – HP1C. (G-I) Identification of DRE motif in promoters bound by HP1a, HP1B and HP1C, respectively.

**Supplemental Figure 2. The genome-wide binding landscape of the Drosophila HP1 family in S2 cells.** (A) Venn diagram illustrating the number of protein-coding genes bound by different combinations of HP1 proteins; Blue – genes bound by HP1a, Green – genes bound by HP1B, Purple – genes bound by HP1C. (B) Proportions of annotated DNA sequence elements (Y-axis) bound by respective HP1 proteins; Blue – HP1a, Green – HP1B, Purple – HP1C. X-axis: sequence element classificiation. (C-E) Proportions of protein-coding genes (Y-axis) bound by HP1a, HP1B, and HP1C in different chromatin contexts (X-axis). (F-H) Comparison of HP1 binding peak width across chromatin contexts (* = p < 1 * 10^-10^, chi-square test); (F) – HP1a; (G) – HP1B; (H) – HP1C. (I) Distribution of protein-coding genes in the Drosophila genome across modENCODE chromatin states. (J-L) Distribution of HP1a, HP1B and HP1C genic binding targets across modENCODE chromatin states.

**Supplemental Figure 3. Increased promoter proximal pausing at HP1 binding targets in S2 cells.** (A-C) Metagene GRO-Seq profiles at HP1a, HP1B, and HP1C binding targets show a higher 5’ peak and greater nascent RNA production at HP1 binding targets. X-axis: scaled position from TSS to TES; Y-axis: RPM. (D-F) Comparison of pausing indices calculated from GRO-Seq data at HP1 binding targets (“Bound”, colored) and non-targets (“Unbound”, grey) in S2 cells; (D) – HP1a; (E) – HP1B; (F) – HP1C. (G) Metagene GRO-Seq profiles of genes when grouped by the combination of HP1 proteins present at the TSS. X-axis: scaled position from TSS to TES; Y-axis: RPM. (H) Pairwise comparisons of pausing indices calculated from GRO-Seq data at genes bound by different combinations of HP1 proteins. X-axis: combination of HP1 proteins present at TSS. Y-axis: Log(Pausing Index). FDR adjusted p values for pairwise comparisons are presented in corresponding heatmap.

**Supplemental Figure 4. Pausing indices across chromatin contexts in third instar larvae.** Metagene profiles of RPII ChIP-Seq data across HP1 binding targets in heterochromatin (A, E, I) and euchromatin (B, F, J) with Y axis showing RPM and X-axis showing the scaled position from TSS to TES. Comparison between pausing indices at HP1 targets (“Bound,” color) and non-targets (“Unbound,” grey) in heterochromatin (C, G, K) and euchromatin (D, H, L). A-D – HP1a; E-H – HP1B; I-L – HP1C. (* = p < 0.0002, permutation test).

**Supplemental Figure 5. Pausing indices across chromatin contexts in S2 cells.** Metagene profiles of GRO-Seq data across HP1 binding targets in heterochromatin (A, E, I) and euchromatin (B, F, J) with Y-axis showing RPM and X-axis showing the scaled position from TSS to TES. Comparison between pausing indices at HP1 targets (“Bound,” color) and non-targets (“Unbound,” grey) in heterochromatin (C, G, K) and euchromatin (D, H, L). A-D – HP1a; E-H – HP1B; I-L – HP1C. (* = p < 0.0002, permutation test).

**Supplemental Figure 6. Histone modification profiles of RPII-elongation related histone marks at HP1 target genes.** HP1a, HP1B, HP1C, H2B-ubiquitination, H3K79me1 and H4K20me1 profiles at HP1 target genes within euchromatin (A), heterochromatin (B), and on chromosome four (C). Genes are classified by the combination of HP1 proteins present at the promoter (left). Only genes that were at least 250 bp away from their nearest neighbor were included for plotting purposes. Color intensity within heatmap reflects coverage at particular loci. Profiles of certain binding combinations of HP1 proteins are excluded due to the small number of genes within that group.

**Supplemental Figure 7. Changes in gene expression in null mutants of respective HP1 family members.** (A-C) Changes in gene expression upon depletion of HP1 proteins. X-axis: Log_2_(Fold Change); Y-axis: negative Log(adjusted p value). Genes without changes in expression are plotted in grey. Genes with significant differential expression are highlighted in color; (A) – differential expression in *Su(Var)205* null mutant larvae. (B) – differential expression in *HP1b* null mutant larvae. (C) – differential expression in *HP1c* null mutant larvae. (D-F) Gene ontology analysis of differentially expressed genes. Y-axis: Fold enrichment (grey) and adjusted p value (color); (D) – HP1a (blue); (E) – HP1B (green); (F) – HP1C (purple). (G-I) Breakdown of overlap between HP1 binding targets and genes differentially expressed upon HP1 depletion; (G) – HP1a (blue); (H) – HP1B (green); (I) – HP1C (purple).

