## Supplementary figures and images for "The Drosophila HP1 family is associated with active gene expression across chromatin contexts"

### Supplemental Figure 1

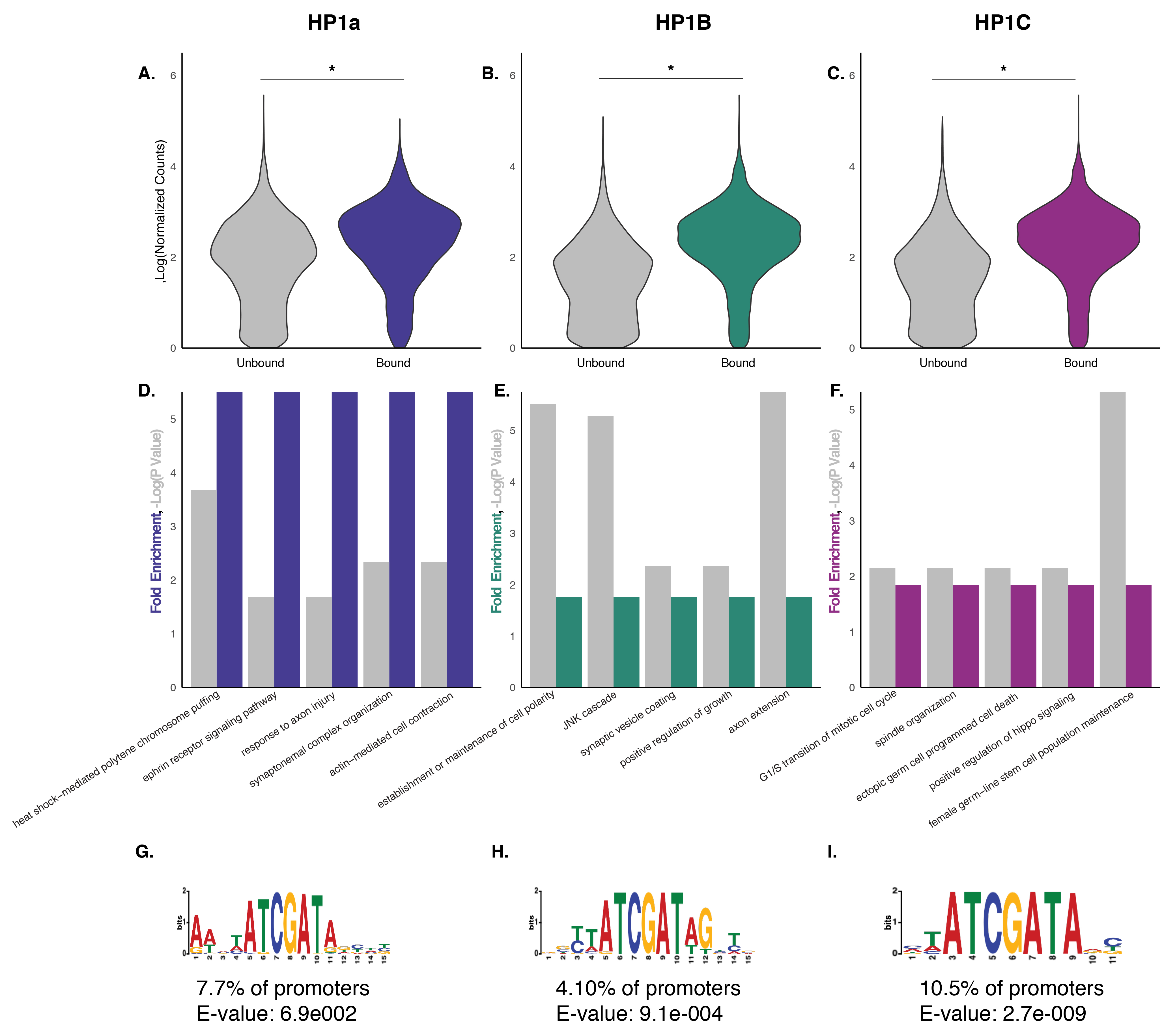

### Supplemental Figure 2

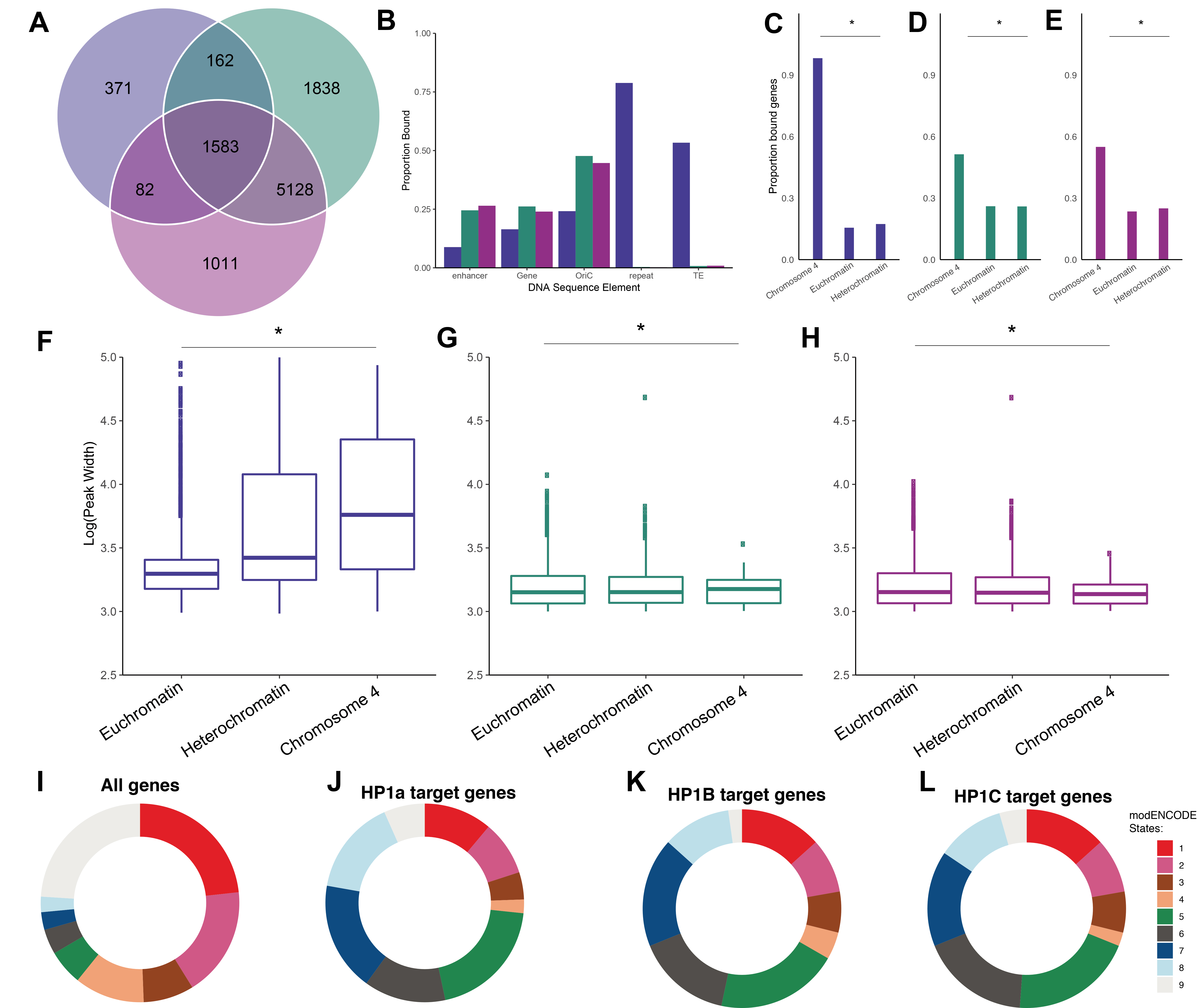

### Supplemental Figure 3

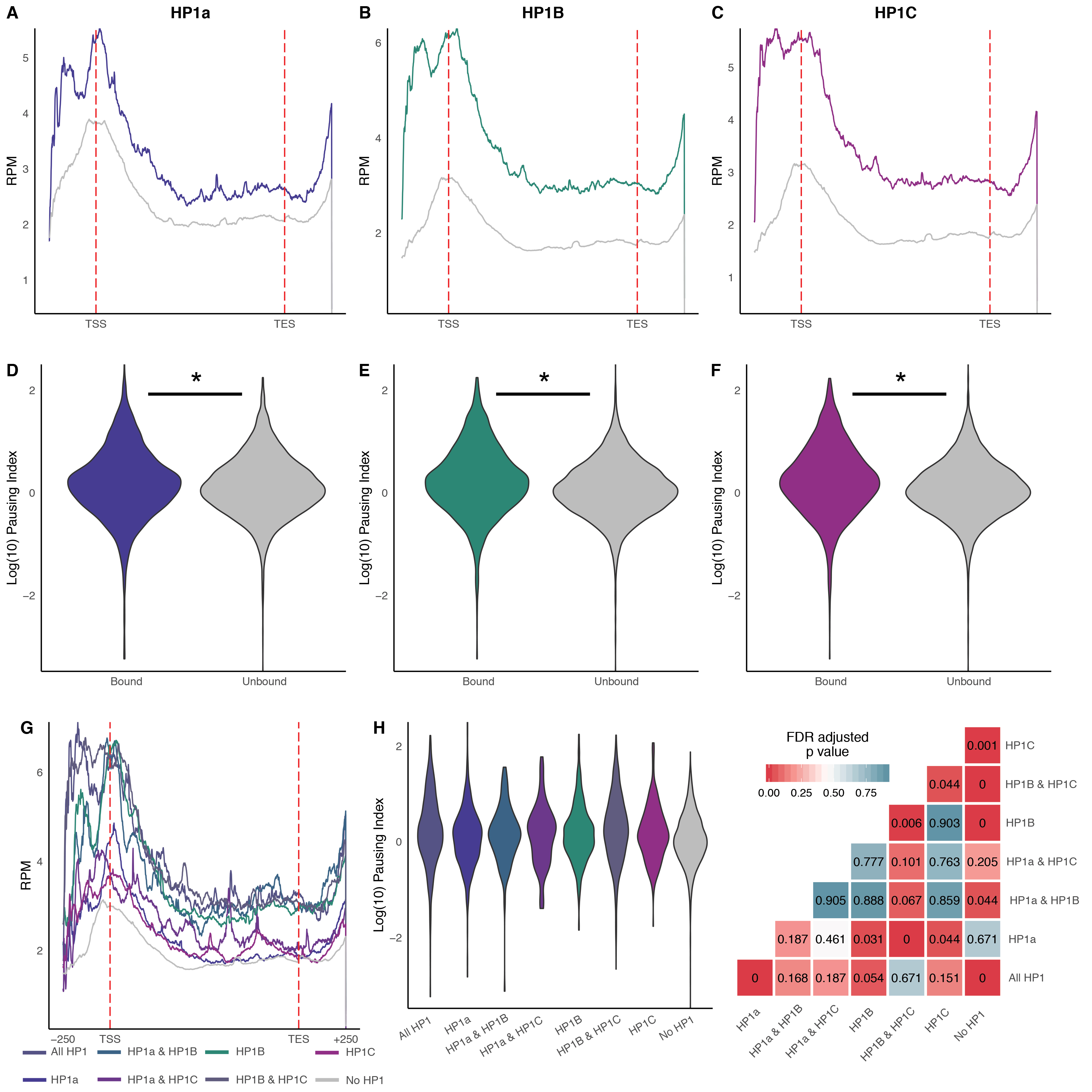

### Supplemental Figure 4

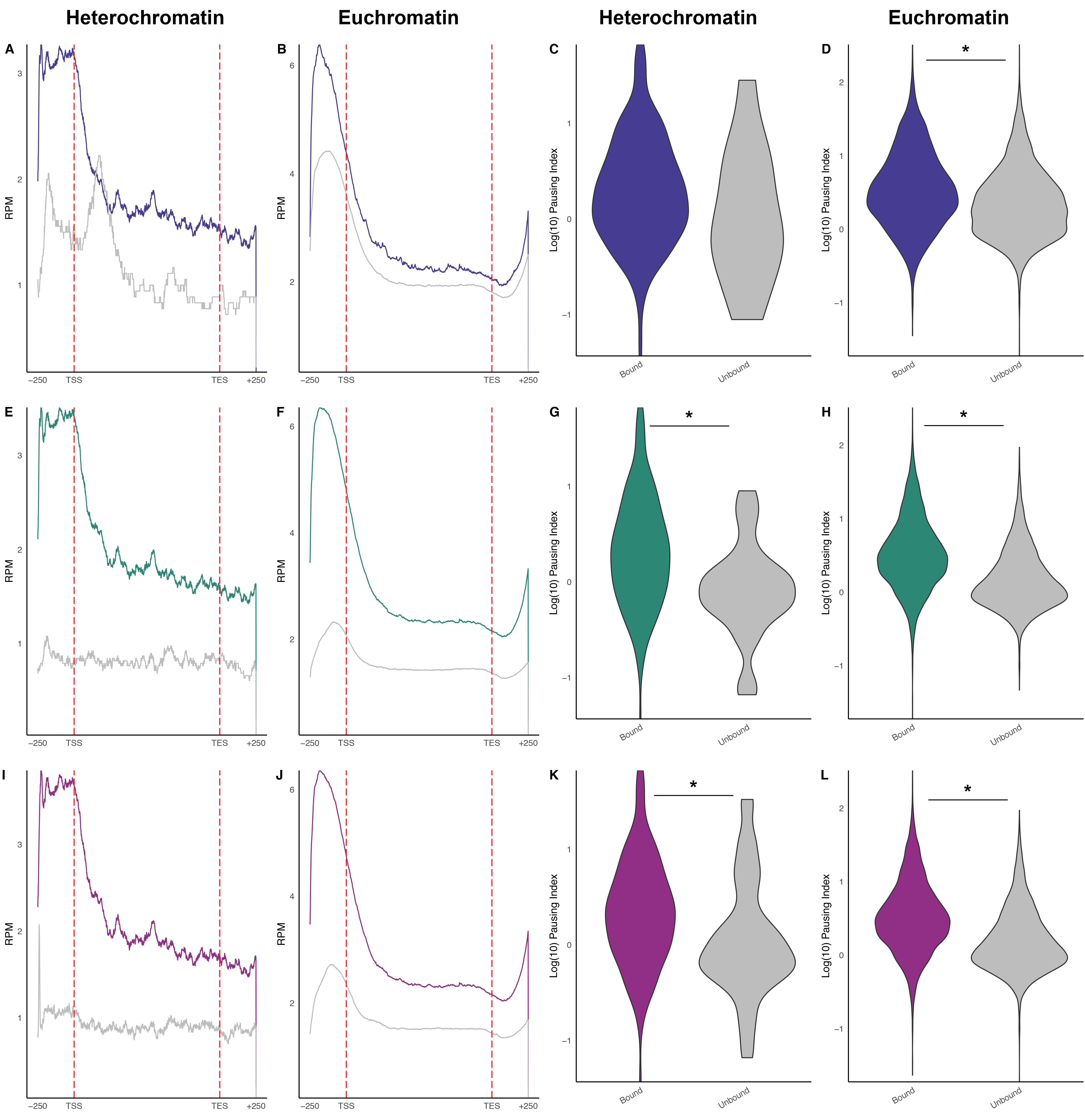

### Supplemental Figure 5

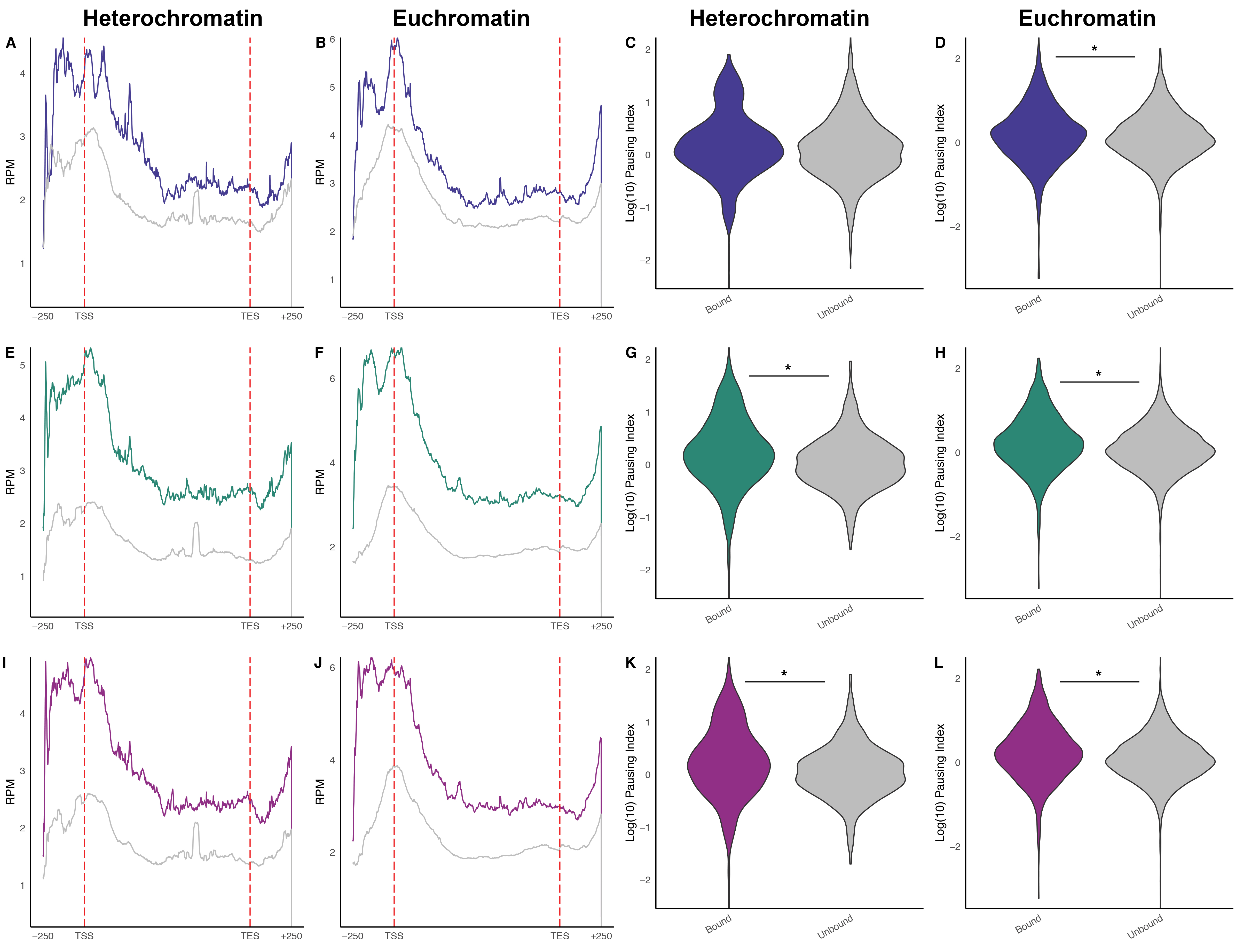

### Supplemental Figure 6

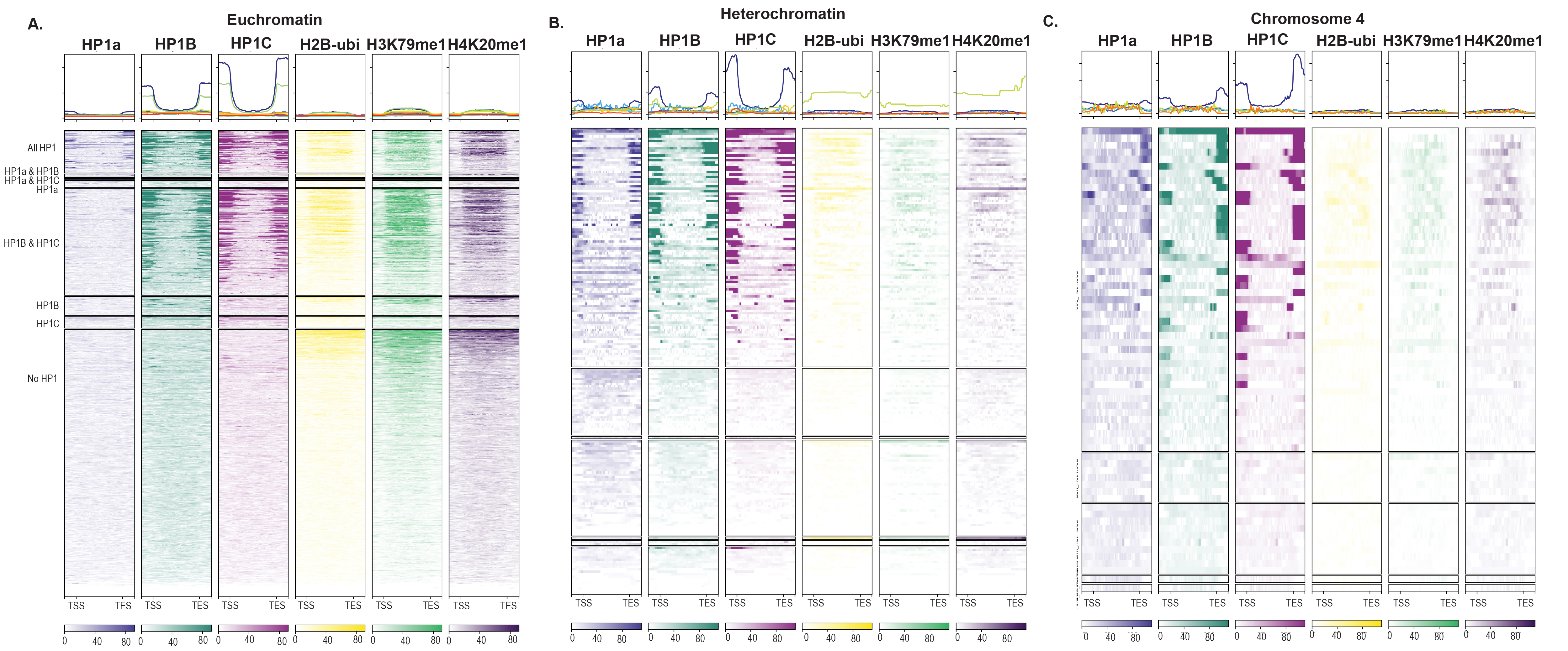

### Supplemental Figure 7

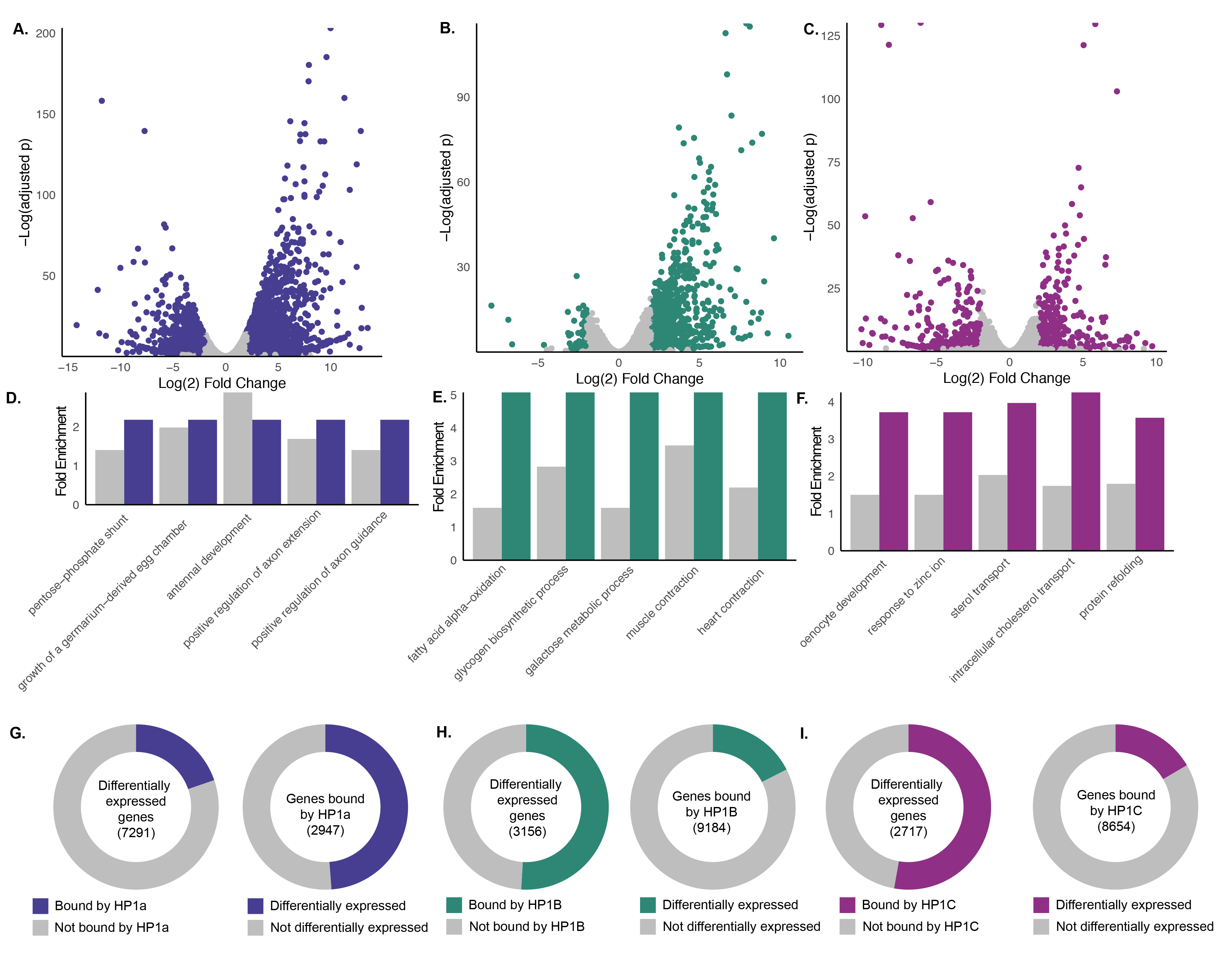
